# The cistrome response to hypoxia in human umbilical vein endothelial cells

**DOI:** 10.1101/2025.03.23.644824

**Authors:** Ayush Singh, Viktor Pastukh, Justin T. Roberts, Zachary M. Turpin, Zehta S. Glover, Grant T. Daly, Jane M. Benoit, Mark N. Gillespie, Hank W. Bass

## Abstract

Hypoxic stress triggers transcriptional signaling mainly through hypoxia-inducible transcription factors (HIFs), which bind hypoxia response elements (HREs) in gene regulatory regions. However, only a small proportion (∼1%) of known HREs are occupied by HIFs during hypoxia, suggesting the involvement of additional hypoxia-responsive factors. To address this gap, we utilized MNase-defined cistrome Occupancy Analysis sequencing (MOA-seq), with the term cistrome referring to all genomic regions where transcription factors and other trans-acting regulators are bound to cis-acting elements across the genome for a particular cell type or treatment. This MNase-based assay enables genome-wide, high-resolution (<30 bp) identification of transcription factor (TF) occupancy footprints embedded within larger regions, most of which were previously annotated as open or accessible chromatin. Applying this *in situ* cistrome mapping to fixed nuclei from endothelial cells under normoxia or hypoxia (1, 3, or 24 hours) revealed thousands of hypoxia-responsive genomic sites with dynamic TF footprints. The affected genes were enriched in canonical hypoxia-induced pathways, such as angiogenesis. Motif analysis identified over 100 candidate TFs potentially mediating these multifaceted genomic responses. By grouping hypoxia-modified occupancy signals across the hypoxia exposure times, we clustered differentially occupied MOA sites into defined 10 distinct TF kinetic clusters, half of which were associated with HIF1A. HIF1A-proximal binding sites suggested co-activators, while non-HIF1A clusters pointed to additional TFs that may have HIF1A-independent roles. This analysis provides insight into how multiple TF networks coordinate hypoxia responses and highlights the power of cistrome profiling to deepen our understanding of the complex genomic response to low oxygen conditions.

**KEY POINTS:**

1. MOA-seq mapped TF occupancy at 21,765 sites in normoxia, including 7,444 beyond the known ENCODE cCREs.
2. Hypoxia for 1, 3, and 24h changes the cistrome occupancy at thousands of genes.
3. Clustering analysis of hypoxia-responsive footprints consolidated cistrome kinetics into HIF1A-associated and HIF1A-independent TFs.

## Introduction

Oxygen availability is a fundamental determinant of cellular function, and in vascular endothelial cells, hypoxia triggers an extensive transcriptional response crucial for angiogenesis, metabolic adaptation, and cell survival. This response is largely orchestrated by hypoxia-inducible factors (HIFs), especially HIF1A, which bind to hypoxia response elements (HREs) and activate key gene networks (Kaluz et al., 2008). Despite extensive study, the determinants of HIF1A binding dynamics across the genome remain poorly understood. Only a small fraction of HREs are actually occupied by HIF1A during hypoxia, and there is limited agreement on which genomic loci are targeted or how co-regulatory transcription factors (TFs) influence binding (Mole et al., 2009; Schödel et al., 2011).

A growing body of evidence suggests that HIF1A does not function in isolation but operates in concert with other transcription factors, chromatin remodelers, and co-activators (Kindrick & Mole, 2020; Villar et al., 2012; Yfantis et al., 2023). These co-regulatory partners influence both the specificity and amplitude of HIF1A-dependent transcriptional programs and help shape the temporal dynamics of gene expression during hypoxic stress. In parallel, hypoxia also engages additional transcriptional and signaling mechanisms that operate independently of HIF1A, particularly in the context of pulmonary hypertension, ischemic injury, and cancer (Pullamsetti et al., 2020; Wu et al., 2021). These include mitochondrial reactive oxygen species (ROS) signaling, which regulates cellular metabolism, apoptosis, and redox balance, and has been implicated in disease progression through the establishment of pseudohypoxic states (Hernansanz-Agustín et al., 2021; Lee et al., 2020; Wu et al., 2021). Additionally, shifts in intracellular sodium and potassium gradients during hypoxia can activate excitation–transcription coupling mechanisms, altering gene expression through ion-sensitive signaling cascades even in the absence of HIF1A (Koltsova et al., 2014). Collectively, these canonical and noncanonical programs contribute to a coordinated transcriptional response that governs how cells adapt to oxygen deprivation. They regulate processes such as metabolic reprogramming, vascular remodeling, proliferation, and survival, with downstream consequences that are tightly context-dependent. In vascular endothelial cells, for instance, these mechanisms help mediate angiogenesis and barrier integrity, whereas in tumor cells, they may promote invasive behavior and resistance to apoptosis (West, 2017). The diversity of hypoxia-induced responses across cell types and tissues underscores the complexity of this regulatory network. Moreover, the hypoxic transcriptional landscape is highly cell-type specific, shaped by epigenetic state, chromatin accessibility, and the availability of cooperative transcription factors, complicating efforts to define universal features of hypoxic gene regulation (Chi et al., 2006). Understanding the upstream genomic events that initiate and coordinate these diverse responses—particularly the shifting patterns of transcription factor occupancy—remains a critical challenge. As such, a comprehensive, time-resolved map of transcription factor activity during hypoxia, one capable of distinguishing canonical HIF1A-driven programs from broader transcriptional dynamics, is essential for elucidating the regulatory logic of hypoxia adaptation and its role in disease pathogenesis.

Traditional methods like chromatin immunoprecipitation sequencing (ChIP-seq) have provided valuable insights into individual transcription factor (TF) binding events, but they offer limited resolution and require high-quality, factor-specific antibodies. Moreover, they often lack the scalability or sensitivity needed to disentangle overlapping TF networks, especially in dynamic environments like hypoxia where multiple transcriptional response programs occur on different but overlapping time scales.

Similarly, methods like assay for transposase-accessible chromatin with sequencing (ATAC-seq) and DNase I hypersensitive sites sequencing (DNase-seq) provide useful information about open chromatin, which generally refers to regions a few hundred base-pairs in length and prone to partial nuclease cleavage within chromatin, but with limited resolution to identify signatures of transcription factors occupancy at specific regulatory elements within these larger open chromatin regions (Yan et al., 2020). As a result, the broader architecture of TF co-occupancy and regulatory network transitions under hypoxia remains poorly understood.

To address this gap, we applied MNase-defined cistrome-Occupancy Analysis (MOA-seq), a high-resolution chromatin footprinting method (Savadel et al., 2021), to human umbilical vein endothelial cells (HUVECs), a model system frequently used to investigate transcriptional responses to hypoxia (Niskanen et al., 2018). The term cistrome is used here to refer to all genomic regions where transcription factors and other trans-acting regulators bind to cis-acting elements across the genome. These comprehensive maps of transcription factor occupancy sites can vary across genotypes or experimental treatment (Engelhorn et al., 2025). Because MOA-seq libraries give genomic data footprints at locations of protein-bound DNA fragments within open chromatin regions, they greatly narrow down candidate TF occupancy to high-resolution, ∼30 bp fragment center peaks (Savadel et al., 2021). Unlike ChIP-seq, MOA-seq is antibody-independent yet comprehensive, enabling precise genome-wide and time-resolved cistrome mapping in a single assay (Engelhorn et al., 2025; Hu et al., 2024; Parvathaneni et al., 2020; Rodgers-Melnick et al., 2016). This approach allowed us to construct time-resolved cistromes in response to hypoxia at 1, 3, and 24 hours for comparative motif analyses.

Our findings revealed a modular and biphasic architecture of transcription factor engagement during hypoxia, driven by both canonical and noncanonical regulators. By expanding the known human cistrome, this study provides a deeper understanding of how hypoxia triggers genome-wide changes in transcriptional regulation and highlights the complexity of hypoxic signaling, including and beyond the well-known HIF1A paradigm.

## Results

### Transcription Factor Occupancy Profiling with MOA-seq in normoxic and hypoxic HUVECs

To investigate hypoxia-induced changes in transcription factor occupancy, we applied MOA-seq to HUVECs under normoxic and hypoxic conditions. For this study, we adapted the assay from maize to human cells using chromatin fixation and MNase digestion protocols previously optimized for mammalian systems (Benoit et al., 2021). This allowed us to generate reproducible, high-resolution cistrome maps across three time points of hypoxic exposure (1, 3, and 24 hours), enabling quantitative analysis of TF occupancy dynamics during the hypoxic response.

The MOA-seq experimental scheme and samples used in this study are summarized in Figure 1. These four samples are designated as 0h, also referred to as normoxia or the pre-treatment control, and 1h, 3h, or 24h to indicate the time of exposure to hypoxia, 2% oxygen in this case. Cells were fixed with formaldehyde to capture the endogenous chromatin structure at the time of fixation. The DNA-protein interactions are revealed as nuclease digestion patterns (Fig. 1B) based on measures of genome-aligned fragment abundances (Supplemental Table S1). This experimental scheme illustrates how small fragment libraries from light nuclease treatments allow for mapping of cis-element occupancy within genomic regions frequently annotated as either accessible or open chromatin. The precision of this footprinting within open chromatin is further increased by fragment center profiling, which focuses the peak regions to ∼ 30 bp, ideal for downstream motif discovery and analysis (bottom of Fig. 1B). Equally important is the fact that these small, high-precision footprints reduce the false-positive error rates that otherwise implicate nearby but unoccupied *cis-*element motifs (see Fig. 1B, cis-element 3) under larger nucleosome-free regions mapped by ATAC or DNAseI peaks.

**Figure 1.**
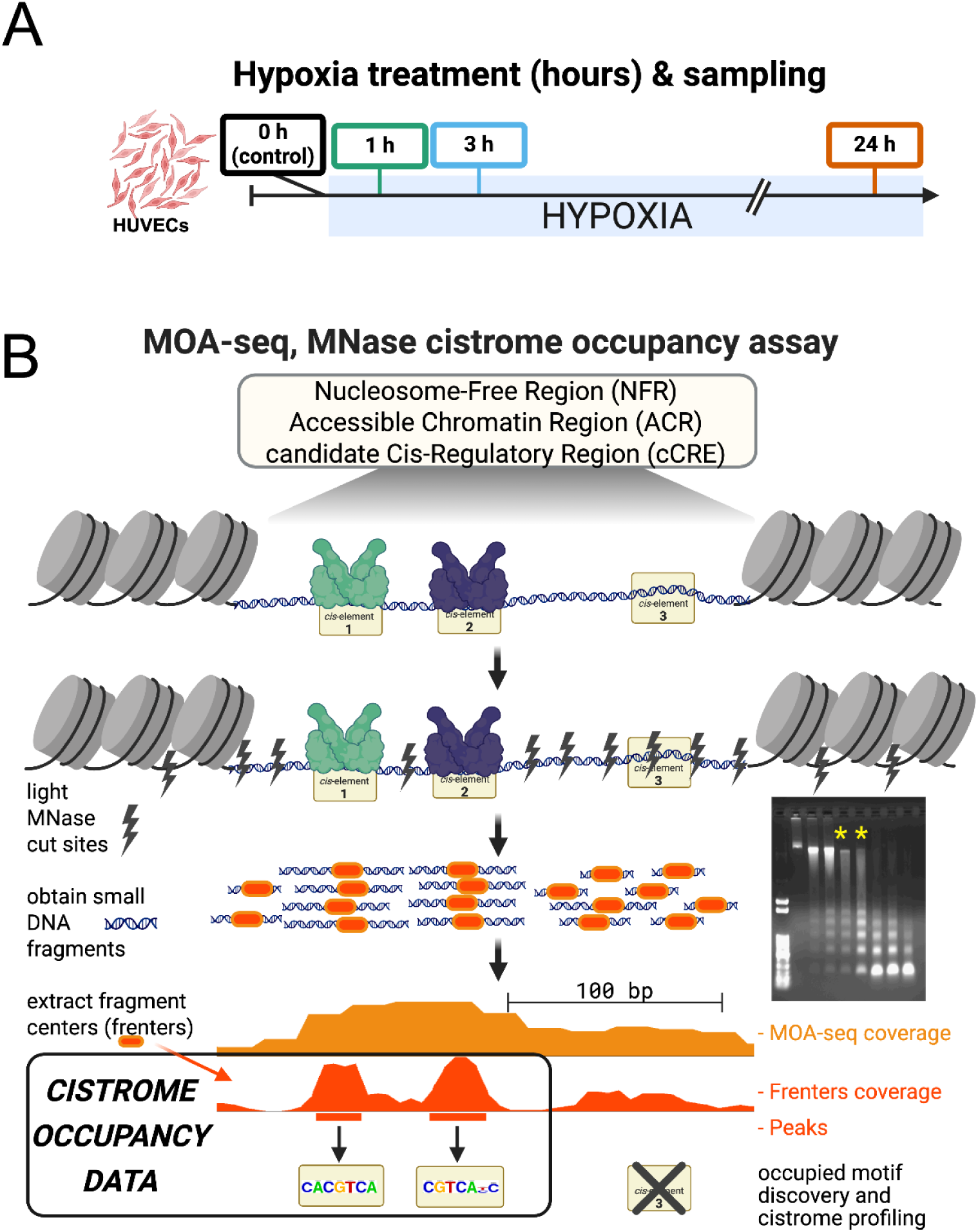
Transcription Factor Occupancy Profiling with MNase "MOA-seq" in HUVECs following hypoxia for 1, 3, and 24h. MOA-seq, <u>M</u>Nase-defined cistrome-<u>O</u>ccupancy <u>A</u>nalysis, (Savadel et al. 2021), was developed to globally identify protein-bound cis-regulatory sequences in a single robust assay using modified nuclease profiling of fixed chromatin and customized bioinformatic analysis pipelines. MOA-seq was applied to HUVEC cells stressed with hypoxia to map short (1h and 3h) or long (24h) exposure-induced changes in TF occupancies. **(A)** The treatment scheme for the four samples used in this study: 0h = normoxia; 1h, 3h, and 24h refer to hypoxia (2% oxygen) exposure times. **(B)** Diagrammatic summary of the MOA-seq, starting with formaldehyde-fixed cells to preserve chromatin structure including bound transcription factors (TFs) at their cognate binding sites (cis-elements). Light/partial digestion with MNase preferentially cuts DNA within nucleosome-free/accessible chromatin regions (NRF/ACR), but not where protein occupancy occurs. Recovery of de-crosslinked, small DNA fragments from ideal partial digests (yellow asterisks, inset gel) is used for NGS library production. MOA-seq coverage profiles of aligned read pair fragments (orange) and their fragment centers (frenters, dark orange) reveal footprints for analysis. The frenter peaks reflect protein-protected footprints within the NFR/ACRs (e.g. cis-element 1 and 2) but do not mark unoccupied cis-elements (e.g. cis-element 3). Peak regions are used for comparative genomics and motif discovery and analysis.

### MOA-seq coverage and footprints align with ENCODE regulatory sites and other genomic regions

We first tested the efficacy of MOA-seq to map sites of transcription factor occupancy by comparing our human MOA-seq data in normoxic HUVECs to established regulatory annotations as summarized in Figure 2. Average aligned-read coverage plots centered on gene features showed characteristic signal patterns of local maxima just upstream of the transcription start site (TSS) and a local depletion within gene bodies (Fig. 2A). At the transcription end site (TES), another local maxima was evident, with a double peak structure flanking the processed transcript end point (Fig. 2B). In both regions, the MOA-seq signals co-directionally scaled with transcript abundance quartiles, indicating that MOA-seq signals generally mirror gene activity, as measured by mRNA levels, while marking canonical regulatory sites with high resolution.

**Figure 2.**
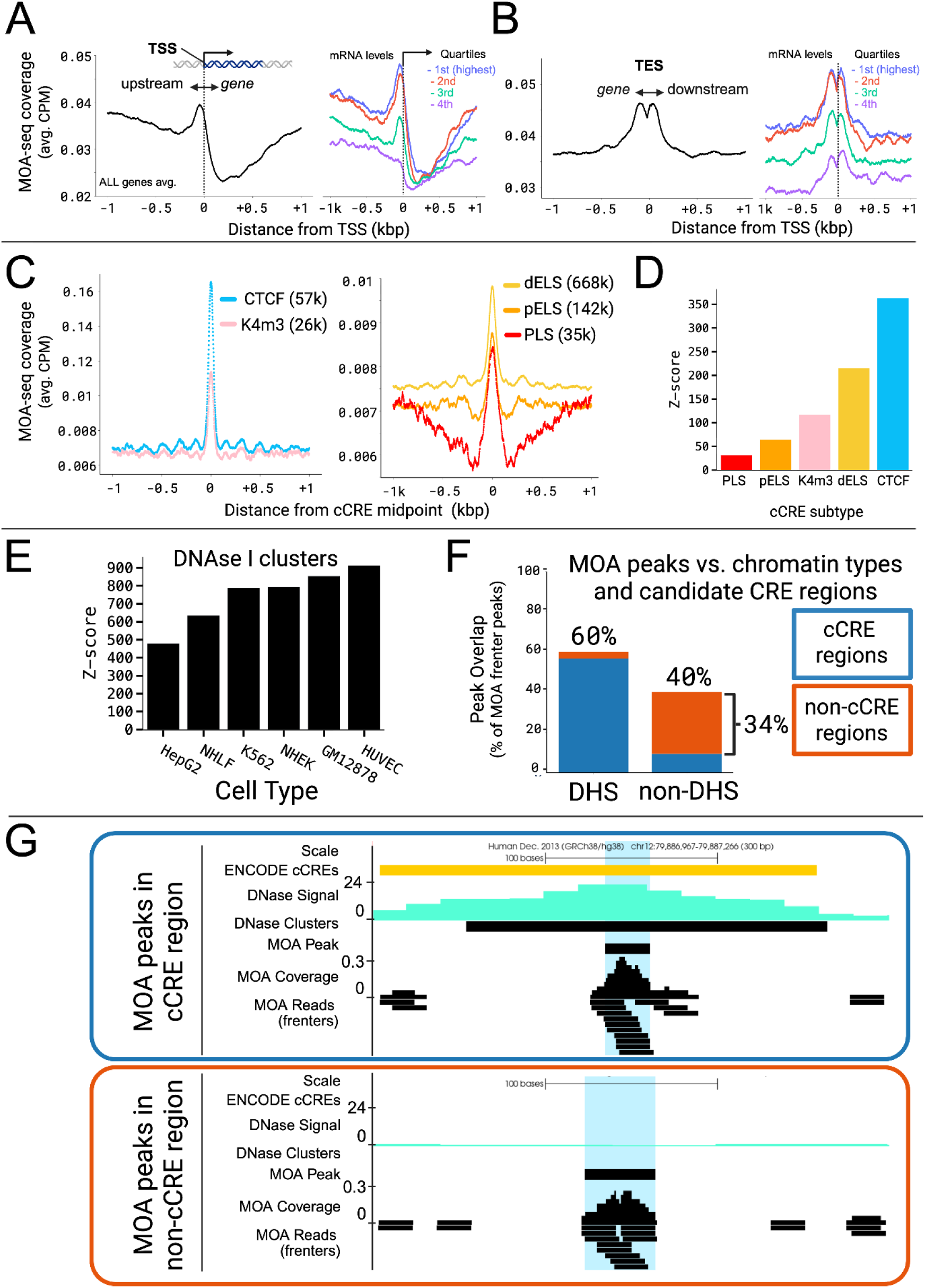
MOA-seq maps gene regulatory features at ENCODE cCREs and additional sites with HUVEC cell-type specificity in normoxic cells. MOA-seq coverage profiles were compared to ENCODE and other genomic annotations related to gene expression and regulation. **(A,B)** Profile trend plots of MOA coverage showing peaks at the transcription start (TSS) and end (TES) sites of genes, with peak heights positively correlating with transcript abundance quartiles. **(C)** Profile trend plots of MOA frenter coverage profiles colocalize with cCRE subtypes (K4m3, DNase I-H3K4me3; dELS, distal enhancer-like signature; pELS, proximal enhancer-like signature; PLS, promoter-like signature). **(D)** Intersection significance is shown by Z-scores calculated from peak bp intersections with cCRE subtypes. **(E)** Intersection significance is shown by Z-scores calculated from peak bp intersections with DNase I hypersensitive (DHS) peaks from multiple different cell types. **(F)** The abundance of MOA-peaks relative to DHS regions or known cCRE regions. **(G)** Genome browser session showing an example of a MOA peak within a cCRE and DHS region (top) and another example in a non-cCRE and non-DHS region (bottom), with tracks labeled on the left.

To better resolve footprint patterns, we restricted peak calling to the central 21 bp of each paired-end fragment, termed “frenters,” which enhances resolution in MNase-based assays (Savadel et al., 2021). Using these frenters, we assessed MOA-seq enrichment at regulatory elements defined by the Encyclopedia of DNA Elements (ENCODE) candidate cis-regulatory element (cCRE) catalog (ENCODE Project Consortium et al., 2020), including the categories promoter-like (PLS), proximal enhancer-like (pELS), distal enhancer-like (dELS), DNase I–H3K4me3 (K4m3), and CCCTC-Binding Factor (CTCF)-only regions. MOA-seq coverage signals showed clear tendencies to colocalize with all five cCRE subtypes (Fig. 2C). The robustness of MOA-seq to detect such localized enrichments was tested by downsampling aligned reads, which ranged from 16M - 41M aligned reads per bioreplicate sample.

Downsampling analysis was carried out for two sets of published reference peaks; the CTCF-only regions cCRE (“An Integrated Encyclopedia of DNA Elements in the Human Genome,” 2012; Landt et al., 2012) peak midpoints and for the ETS Transcription Factor ERG (ERG) motif under the ERG ReMap ChIP-seq peaks. ReMap is a curated database of human transcriptional regulator binding peaks derived from ChIP-seq, ChIP-exo, and DAP-seq experiments (Hammal et al., 2022). We found that the MOA coverage signal was essentially the same even after downsampling to 1M reads (Fig. S1), indicating a sequencing depth of well over ten fold greater than required.

Using the Irreproducibility Discovery Rate (IDR) framework, which identifies reproducible peaks across biological replicates ((Q. Li et al., 2011), we identified 21,765 high-confidence MOA peaks in normoxic (0h) HUVECs (Supplementary Table S2). Enrichment analysis showed that MOA-seq peaks were highly overrepresented at distal enhancers and CTCF sites (Z-scores > 200; Fig. 2D, Supplementary Table S3), and showed strong concordance with HUVEC DNase I hypersensitivity data (Fig. 2E).

These analyses demonstrate that human cell culture (HUVEC) MOA-seq captured active regulatory elements with high specificity and overlapped open chromatin regions annotated by orthogonal datasets such as DNAse I hypersensitive sites. Approximately 40% of MOA peaks did not overlap DHS clusters, and 34% did not overlap cCREs (Fig. 2F), suggesting that MOA-seq also detects regulatory regions not captured by traditional assays. Representative genome browser tracks further illustrate MOA-seq peaks at both annotated and novel loci (Fig. 2G).

Finally, to characterize the genomic distribution of normoxic MOA peaks, we annotated all 21,765 peaks using ChIPseeker. Only 3.0% mapped to promoter regions (±200 bp from the TSS). The majority localized to intronic (17.2% first intron, 39.8% other introns) and distal intergenic regions (33.5%). These distributions remained consistent among peaks that did not overlap DHSs, cCREs, or either (Fig. S2).

### Enrichment of normoxic MOA at ReMap ChIP summits for transcription factors from HUVEC-specific datasets

To next assess whether MOA-seq signal corresponds to known transcription factor binding sites, we analyzed ChIP-seq summits from the ReMap database (Hammal et al., 2022), focusing on 13 TFs with HUVEC-specific data and high expression in endothelial cells (Payne et al., 2024). As shown in Figure 3, MOA-seq coverage in normoxic HUVECs showed strong enrichment at ReMap ChIP-seq summits (Fig. 3A, B). Individual TF enrichment profiles revealed colocalization of MOA-seq coverage, with RAD21 Cohesin Complex Component (RAD21) and Myocyte Enhancer Factor 2C (MEF2C) showing the highest local enrichment among the 13 HUVEC TFs (Fig. 3C, Fig. S3). These results show that MOA-seq captures binding events for multiple TFs known to be active in endothelial cells.

**Figure 3.**
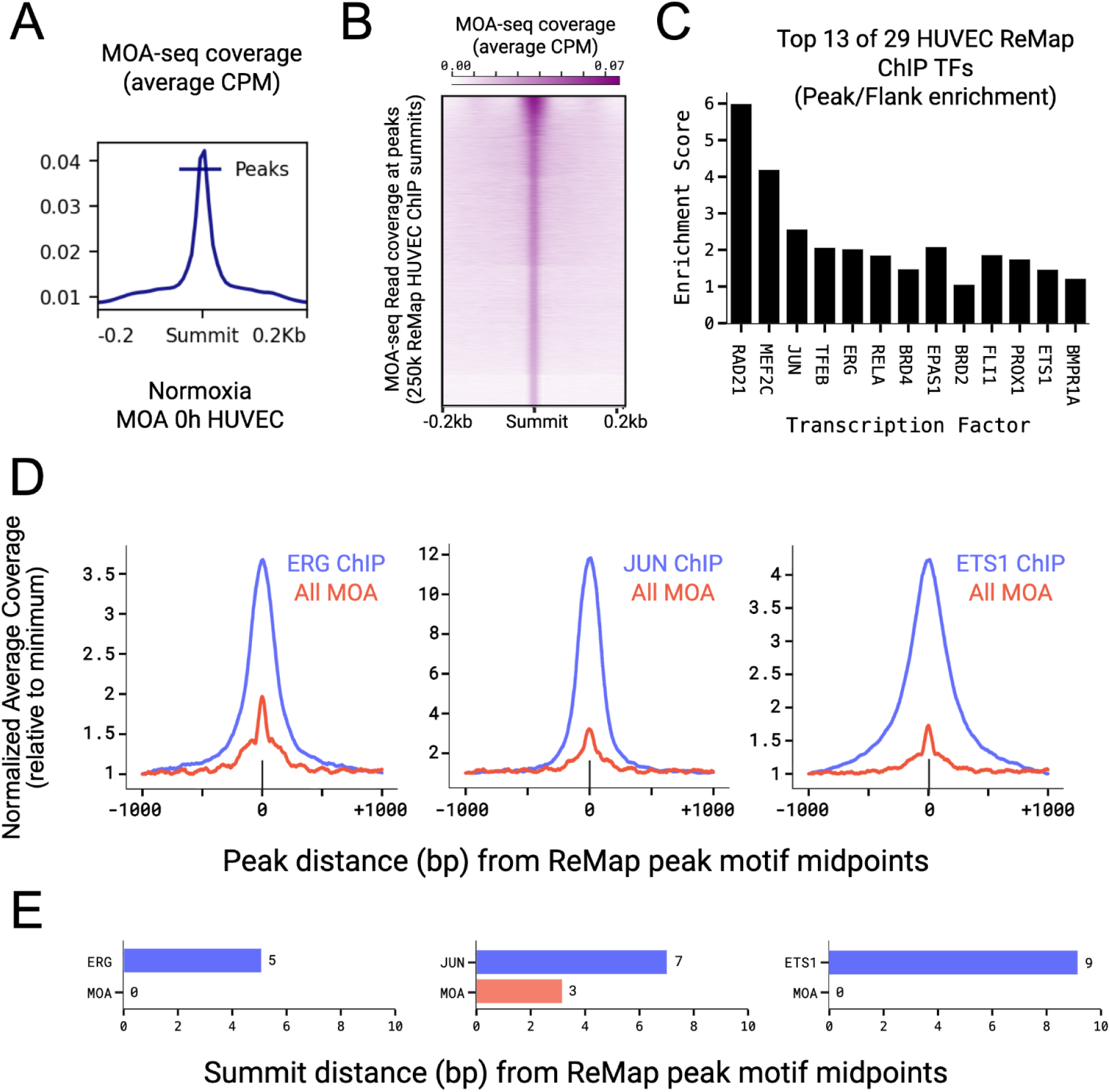
MOA-seq Enrichment at ReMap ChIP Summits for Canonical HUVEC Transcription Factors. MOA-seq enrichment at previously characterized TF binding sites as defined by ChIP-seq, using ReMAP (Hammal et al., 2022) ChIP summits for 13 canonical HUVEC TFs (Payne et al., 2024). Profile trend **(A)** and heatmap **(B)** plots of MOA 0hr frenters coverage centered on ReMap HUVEC ChIP peak summits. Data plotted were from regions with non-zero MOA coverage of ∼250k summits from a total of ∼750k spanning various studies and treatments. **(C)** Bar chart showing total unfiltered enrichment scores (MOA coverage ratio of summit value/flanks, see Methods) for a select subset of TF ChIPs from experiments conducted in HUVEC only, with nearby (< 10 bp) summits for MOA and ChIP. **(D)** Profile trend plots depicting local average reads per million for MOA frenters (red) and ChIP-seq (blue). ChIP-seq data is from individual HUVEC experiments (S. Wang et al., 2019) centered on the canonical motif for the respective transcription factor (TFs). **(E)** Bar charts illustrating the distance (bp) of MOA and ChIP summits from the motif midpoint for each TF.

The ReMap database integrates ChIP-seq data across studies to define TF-specific consensus regions. However, many ReMap peaks overlap with nearby binding sites for multiple other TFs. Therefore, we wanted to explore the degree to which the MOA frenters exhibited precise alignment at the TF motif. We identified canonical binding motifs for ETS Proto-Oncogene 1, Transcription Factor (ETS1), ERG, and Jun Proto-Oncogene, AP-1 Transcription Factor Subunit (JUN) within their respective ChIP-seq peaks and plotted both MOA-seq and ChIP-seq coverage centered on motif midpoints (Fig. 3D). Both signals were concentrated at the motif center, with MOA-seq showing a sharper spike—particularly for ERG and ETS1. To better quantify the positional precision of ChIP versus MOA for select motifs, we calculated the distance between motif midpoints and the coverage summits for ChIP or MOA (Fig. 3E). For ERG and ETS1, the MOA-seq summit was centered on the motif, while the ChIP-seq summit was offset by 5 bp and 9 bp, respectively. For JUN, the MOA-seq summit was 3 bp from the motif center and the ChIP-seq summit 7 bp away. This analysis supports the interpretation that MOA-seq can resolve transcription factor binding at nucleotide-level precision, helping distinguish individual TF occupancy within slightly larger regions defined by ChIP-seq peaks.

### Hypoxia causes temporally-specific changes in MOA-defined chromatin structure

Having demonstrated that MOA-seq is highly informative for mapping TF occupancy within human cell cultures, we next used it to define genomic regions with differential MOA coverage following hypoxia. These regions, designated herein as “diff-MOA” sites, are summarized in Figure 4. We identified diff-MOA peaks using the bdgdiff function in MACS3, optimizing the signal-to-noise ratio by applying the elbow method defining a robust set of MOA peaks that dynamically respond to hypoxia across time points. For each hypoxia time point, we identified three categories of peaks relative to 0h normoxia: gained, lost, or shared. The total number of gain and loss peaks was 1,978 after 1 hour, 1,932 after 3 hours, and 2,586 after 24 hours of hypoxia (Fig. 4A). Global patterns are visualized in heatmaps sorted by MOA-seq signal for gain and loss peaks (Fig. S4).

**Figure 4.**
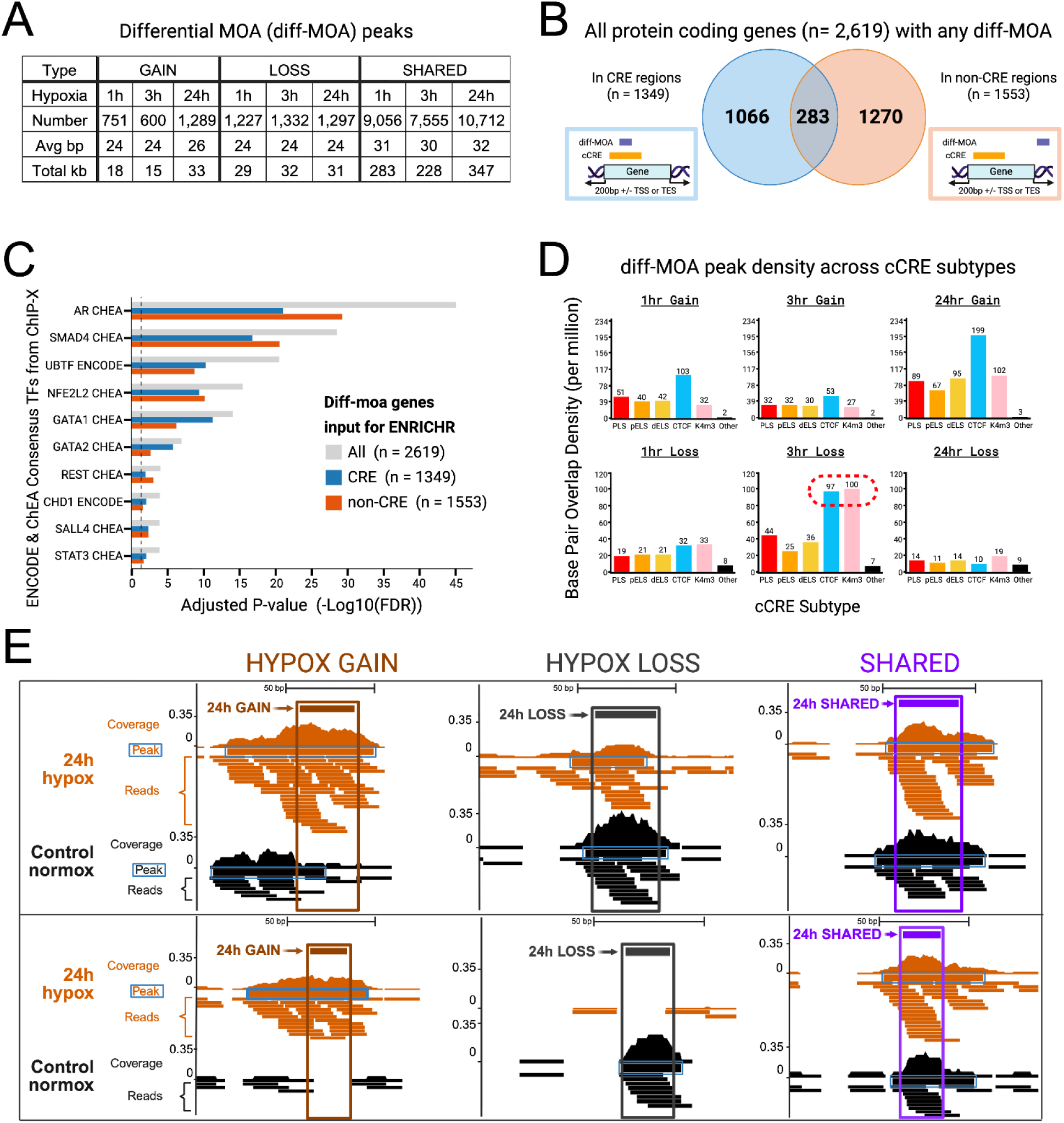
Hypoxia-induced differential MOA (diff-MOA) peak analysis and transcription factor enrichment across genomic regions. To identify genomic regions that changed their TF occupancy footprints in response to hypoxia, bdgdiff (MACS3) of MOA frenter coverage was used to call peaks using the elbow method. **(A)** Table of diff-MOA peak numbers and sizes for diff-MOA peaks. Each is one of the three hypoxia time points relative to 0h normoxia. **(B)** Venn diagram depicting the overlap of genes with diff-MOA footprints in cCRE versus non-cCRE regions. Genes in the intersection contain multiple diff-MOA footprints, with at least one footprint located in a cCRE and one located in a non-cCRE region. Schematics depict genes with a diff-MOA peak overlapping a cCRE region (light blue box) or a non-CRE region (light orange box). **(C)** Enrichments (adjusted p-values) for gene lists derived from diff-MOA peaks in cCRE versus non-cCRE regions. Results from Enrichr (https://maayanlab.cloud/Enrichr/) using the libraries for ENCODE and ChEA Consensus TFs from ChIP-X. The plot was sorted by P-values for the total diff-MOA peaks (grey bars) and all significant TFs (dashed lines) are shown. **(D)** Base pair density overlap (per megabase) of gain and loss peaks across ENCODE cCRE subclasses: K4m3 (DNase I–H3K4me3), dELS (distal enhancer-like signature), pELS (proximal enhancer-like signature), and PLS (promoter-like signature), along with an additional category labeled “Other” representing peaks not overlapping any cCRE subclass. **(E)** Genome browser session showing two examples each for diff-MOA gain, diff-MOA loss, or shared for 24h hypoxia treatment. The within-sample, non-differential peaks are indicated (Peak, blue boxes). The differential, diff-MOA, peaks are shown above each pair, and boxed with a unique color and indicated as 24h GAIN (dark orange), 24h LOSS (dark grey), or 24h SHARED (purple).

We next identified ’diff-MOA genes’—protein-coding genes with a diff-MOA peak located within the gene body, 200 bp upstream of the TSS, or 200 bp downstream of the TES, resulting in a total of 2,619 genes from all time points (Fig. 4B). A substantial fraction of these genes contained diff-MOA peaks that occurred outside of known cCRE annotations (Fig. 4B; Supplementary Table S4). To test for the biological significance of the genes with non-CRE diff-MOA peaks, we performed transcription factor target gene enrichment analysis using the Enrichr platform with ENCODE/ChEA consensus TF target gene sets derived from ChIP-X datasets (Chen et al., 2013; Kuleshov et al., 2016; Xie et al., 2021) on the three groups of gene sets; all 2,619 diff-MOA genes, the 1,349 diff-MOA genes with peaks in CRE regions, and the 1,553 diff-MOA genes without CRE-overlapping diff-MOA peaks (Fig. 4C; Supplementary Table S5). Both subgroups, cCRE-intersecting (n=1,349 genes) and non-cCRE (n=1,553 genes), showed comparable enrichment for known transcription factor target gene sets from ENCODE and CHEA ChIP-X datasets. Randomized gene lists of equivalent size did not show any significant enrichment (adjusted P > 0.05; Supplementary Table S5). We found that diff-MOA peaks, whether in cCREs or not, identified genes with comparable enrichment values for TF gene ontologies, and can be pooled for biological analysis.

We next aimed to determine whether the peaks that were gained, lost, or shared across the three time points are predominantly located within particular types of regulatory regions, such as promoters or enhancers (Fig. 4D; Fig. S5 and S6). We used the ENCODE cCREs which are subdivided into five subclasses, including promoter-like and enhancer-like categories. Gain peaks consistently showed the strongest enrichment at CTCF sites, followed by promoter-like (PLS) and enhancer-like elements (pELS, dELS), except for 24-hour gain peaks, which also exhibited comparable enrichment at DNase I–H3K4me3 sites. In contrast, loss peaks at 1 and 3 hours were primarily enriched (red dashed lines, Fig. 4D) at DNase I–H3K4me3 sites, followed by CTCF sites, neither of which are at promoters. By 24 hours, loss peaks were fewer and more uniformly distributed across the five cCRE categories. Shared peaks followed a similar pattern, with the highest enrichment at CTCF, DNase I–H3K4me3, and distal enhancer sites (Fig. S6). Representative regions illustrating hypoxia-induced changes in chromatin structure are shown in browser snapshots (Fig. 4E). These analyses show that hypoxia causes small footprint changes, both gained and lost, at regulatory sites in thousands of genes.

### Comparison of differential MOA with differential gene expression

We next addressed whether time-dependent changes in the hypoxic transcriptome were biologically related to MOA-defined cistrome dynamics. To do this, we compared two sets of genes based on their response to hypoxia, those harboring diff-MOA peaks and those classified as differentially expressed genes (DEGs) as summarized in Figure 5. Compared to normoxia (the 0h sample), we found 1130, 1092, and 1401, diff-MOA genes at 1h, 3h, and 24h, respectively. Similarly, we found 853, 1,830, and 2,521 DEGs at 1h, 3h, and 24h of hypoxia, respectively.

**Figure 5.**
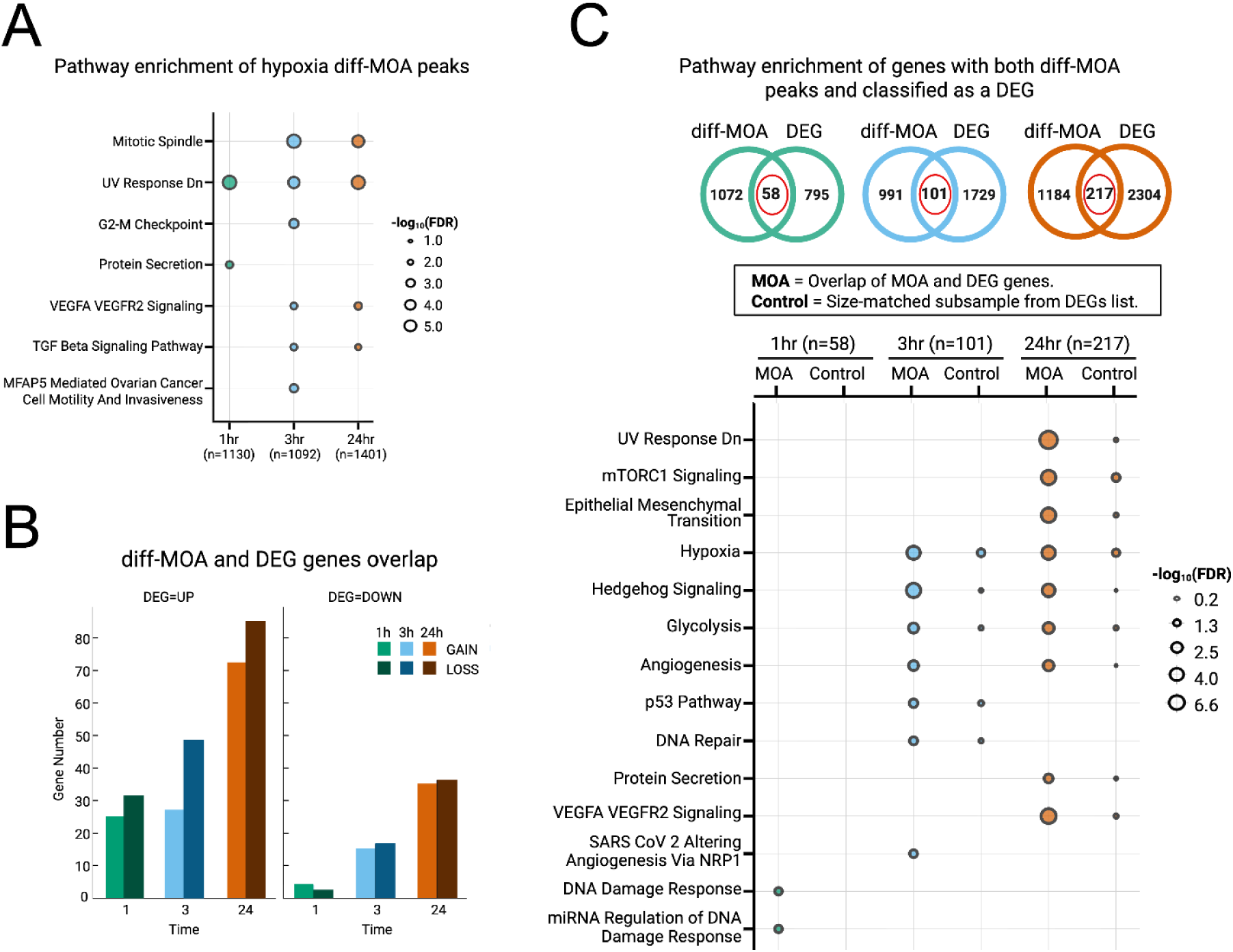
Comparison of differential MOA with differential gene expression. Comparison of hypoxia-induced diff-MOA genes to hypoxia-induced differentially expressed genes (DEGs) with regard to gene enrichment analyses, changes in RNA levels, and gain or loss of diff-MOA footprints. **(A)** Gene lists defined by diff-MOA (sum of all gain- or loss-peak genes) for each hypoxia time point were pooled and used as input gene lists for the Enrichr libraries. Top hits are shown for two libraries, Human MSigDB Hallmark and WikiPathway Human gene sets. The significance value (-log10(FDR)) is proportional to the circle size; gene list sizes are indicated at the bottom. **(B)** Lack of directional relationship between diff-MOA (gain or loss) and DEG (transcript up or down) is shown as gene numbers plotted for each time point. **(C)** Analysis of genes showing both diff-MOA and DEGs. The top shows Venn diagrams for each time point. The bottom shows Enrichr results using the same two libraries as in panel A. Genes with both diff-MOA and DEG gene subsets (MOA) have stronger enrichments as shown by comparative analysis using randomly sampled subset of genes from the list of DEGs (Control). For each time point the average enrichment is plotted based on three size-matched subsets of genes (sampling 58 DEG genes for 1h, 101 DEG genes for 3h, and 217 DEG genes for 24h. For each such comparison, the average (n=3) subsamples significance (-log10(FDR)) was less than for the MOA plus DEG group.

Gene sets defined by diff-MOA peaks alone showed significant enrichment in the Enrichr Human MSigDB Hallmark and WikiPathway Human libraries (Fig. 5A), including the well-characterized hypoxia-responsive "VEGFA–VEGFR2 signaling" pathway, which reached adjusted P-values of 0.034 at 3 hours and 0.015 at 24 hours (Supplementary Table S6). To further explore this relationship, we classified DEGs as up- or down-regulated and diff-MOA genes as gain or loss of hypoxia-induced footprint changes. Only a moderate number of genes overlapped between the two lists. Among these doubly differential genes, there was a slight tendency toward more loss than gain of MOA-defined footprints (Fig. 5B; Supplementary Table S7). However, neither the gains nor losses of footprints were associated with up versus down regulation as measured by DEGs.

To assess the functional relevance genes on both lists, DEGs and diff-MOA, we compared gene ontology (GO) enrichments between doubly differential genes (58 genes for 1hr, 101 genes for 3hr, and 217 genes for 24hr) and size-matched randomized subsets of DEGs for each time point. Notably, gene sets defined by the intersection of diff-MOA and DEG lists showed stronger pathway enrichment than the size-matched DEGs alone, using the same MSigDB Hallmark and WikiPathway libraries (Fig. 5C; Supplementary Table S7). This enhanced enrichment indicates that the intersection of cistrome remodeling and gene expression changes captures a stronger signal of biological relevance for these hypoxia-responsive genes.

### Motif discovery and analysis at diff-MOA peaks nominate *cis*-regulatory elements involved in hypoxia-responsive gene regulatory networks

Because diff-MOA peaks are small, averaging 24 to 32 base pairs (Fig. 4A), and likely reflect TF occupancy (Fig. 3), their underlying sequences are ideal for motif analysis. We divided the peaks into two groups, gain and loss, and submitted their genomic sequences to the MEME Suite for motif discovery. The resulting output represents a very large dataset, typical for this cistrome occupancy assay and akin to pooling thousands of ChIP-seq experiments (Supplementary Zip File 1). The top motifs identified for each group and hypoxia time point are shown in Figure 6, separated into gain-associated motifs (Fig. 6A) and loss-associated motifs (Fig. 6B). This comprehensive analysis yielded six motif sets, one gain and one loss set for each of the three hypoxia time points. Within each set, significant *de novo* motifs are listed first, followed by common TF motif families such as CTCF, Fos Proto-Oncogene, AP-1 Transcription Factor Subunit (FOS), and JUN, and then the top three TF-specific motifs ranked by E-value (Full motif lists in Supplementary Table S8). The third column shows the percentage of diff-MOA peaks that contain each motif, ranging from 2% for Zinc Finger Protein 502 (ZNF502) to 55% for SMAD Family Member 3 (SMAD3). Given that many TFs share similar or overlapping consensus sequences, we also reported the proportion of motifs that intersect with corresponding ChIP-seq peaks from the ReMap database. Motifs from major TF families, including JUN, FOS, and CTCF, were highly prevalent at gain sites at 1 and 24 hours but were absent from loss sites at all time points, except for 3-hour loss peaks, which retained CTCF motifs. This observation is suggestive of a variation in motif composition and TF involvement across time points, revealing substantial complexity of potential regulatory dynamics between gain and loss peak regions.

**Figure 6.**
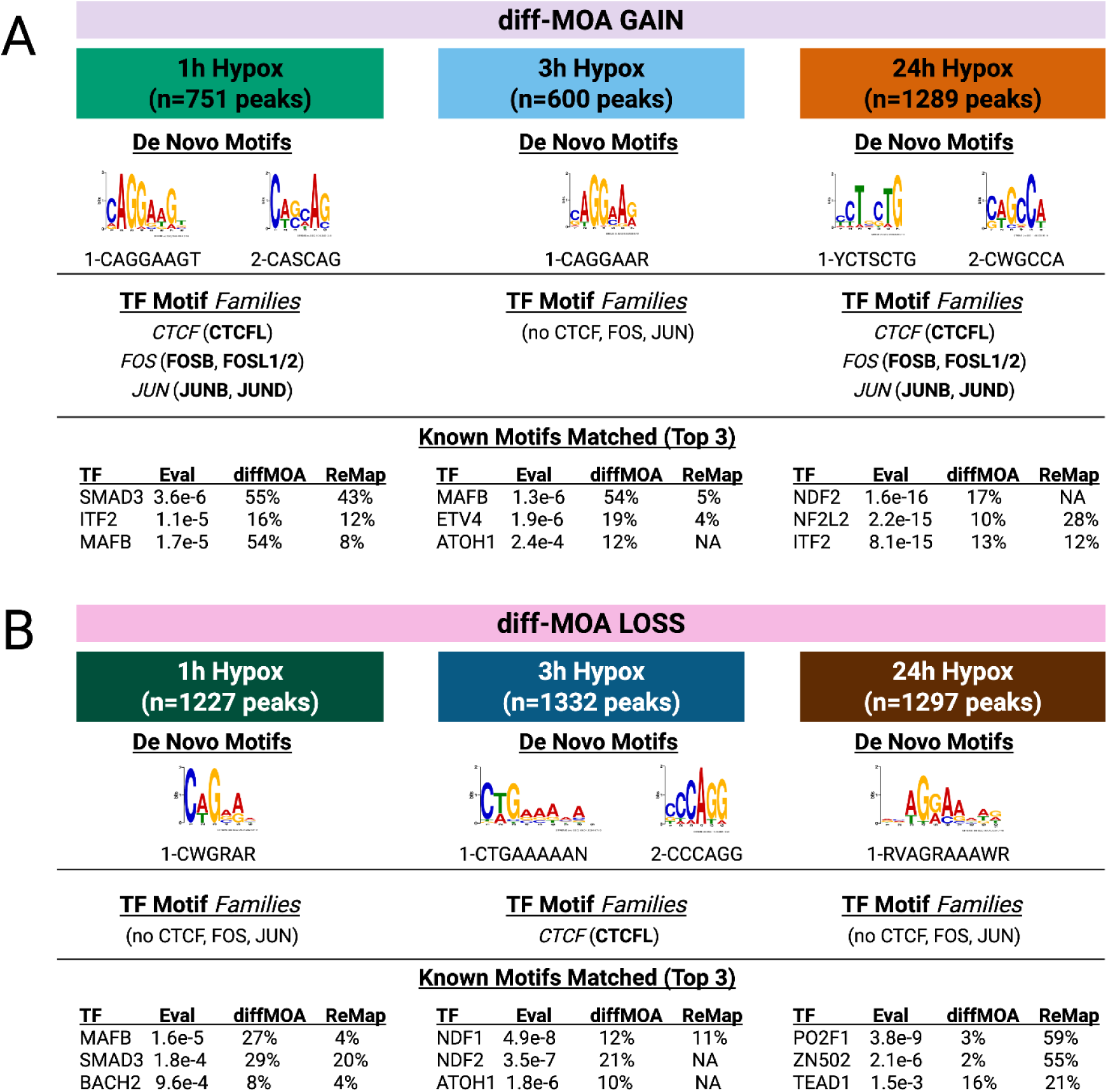
Motif discovery and analysis at diff-MOA peaks identify candidate hypoxia-responsive gene regulatory network cis-elements. The diff-MOA peaks were split into six sets based on gain **(A)** or loss **(B)** of footprints at each of the three hypoxia treatment timepoints. Genomic sequences under the peaks were analyzed using MEME motif analysis. For each list of motifs, three results are shown: the significant de novo motifs with their sequence LOGO, the TF families (italicized) and their members (bolded in parentheses), and the top 3 individual TFs sorted by E-value. For each motif (TF), the background-adjusted expected number of motifs (Eval) is listed in the second column, followed by the percentage of diff-MOA peaks containing the motif (diff-MOA), and the percentage of the motif intersected with ReMap ChIP-seq data (ReMap, NA represents absence of a ReMap ChIP-seq data).

### Clustering MOA coverage kinetics resolves transcription factor motifs and associated functional pathways

To better understand the complexity and scale of the observed changes and to explore how TF networks might drive these diverse responses, we performed a quantitative footprint clustering analysis. This allowed us to capture shared and distinct coverage patterns across time points and conditions based on 5,383 hypoxia-induced regions.

These regions were set at 30-bp windows centered on all diff-MOA peaks as input. We applied an unsupervised clustering approach using k-means clustering with the elbow method (Supplementary Table S9), defining 10 hypoxia-induced MOA-seq clusters, named hmc1 to hmc10 (Figure 7). Some clusters were enriched for early TF binding events (e.g., hmc2 at 1h and hmc4 at 3h), while others showed later-stage occupancy patterns (e.g., hmc3 and hmc9 at 24h). The number of diff-MOA peak regions in the clusters ranged from 64 for hmc10 to 967 for hmc8, with most clusters having over 400 regions (Supplementary Table S10). Each cluster was individually used as input for MEME motif analysis, and the top motifs for each cluster are listed under their respective panels (Fig. 7A, Supplementary Zip File 2). Nearly all the motifs from the genome-wide analysis were retained but conveniently partitioned into one or more clusters (Supplementary Table S11). The clustering improved resolution of the motif families, successfully subdividing them and identifying top TF lists per cluster. For example, the ETS Variant Transcription Factor 4 (ETV4) motif was prominent in hmc1, hmc3, and hmc8, while CCCTC-Binding Factor Like (CTCFL) was primarily limited to hmc10 and hmc2.

**Figure 7.**
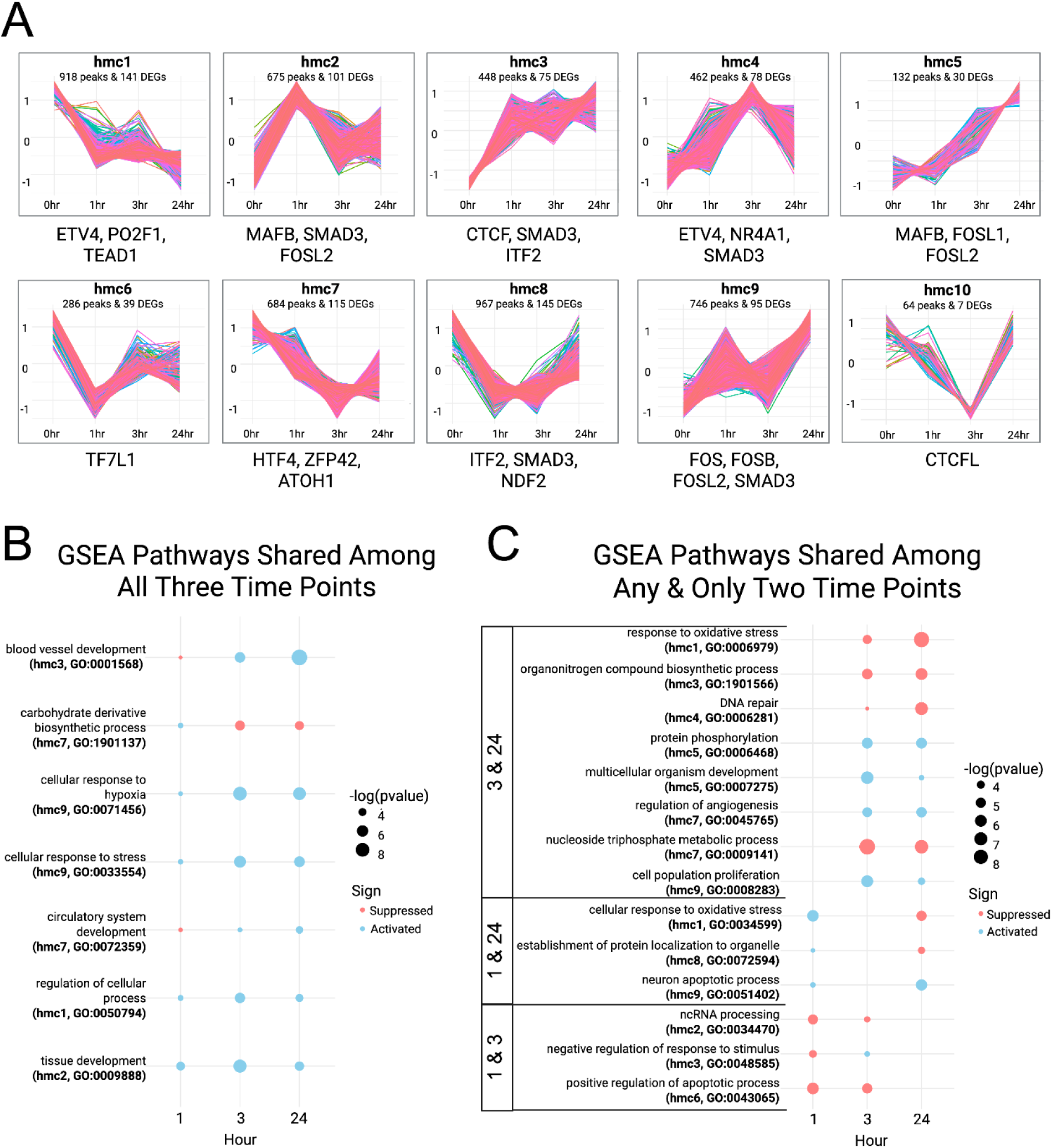
Cluster analysis and GSEA GO enrichment results. We defined 5,383 hypoxia-induced peak regions as 30-bp windows centered on all diff-MOA peaks. The R-package k-means clustering of the MOA coverage values with the elbow method resulted in 10 clusters. **(A)** The <u>h</u>ypoxia <u>M</u>OA <u>c</u>lusters were named "hmc1" through "hmc10", plotted for each peak region using Z-score standardization (Y-axes -1 to +1), along with the total number of peaks and DEGs associated with each cluster. The peak regions for each cluster were separately used as input for MEME motif analysis, and the top motifs for each are listed under their respective panels. **(B, C)** The cluster-specific DEGs and their fold change transcript abundance for each hypoxia treatment hour were used as input for the gene set enrichment program, gseGO in clusterProfiler. Resulting enrichments are shown and plotted across the time points for significant pathways, for suppressed (pink) or activated (blue) genes. Plots list the GO term, the hmc cluster number, and the GO ID for each pathway for those found to be shared among all three time points (left) or shared between any two of the three time points (right).

Because distinct motifs appeared in different clusters, we asked whether the associated genes reflected distinct biological responses. Of the 5,383 diff-MOA regions, nearly 60% overlapped annotated gene regions, which we define as the gene body +/- 200 base pairs to capture promoter and 3’ regulatory regions. We performed Gene Set Enrichment Analysis (GSEA) using cluster-specific gene sets and their fold-change values from our expression data relative to time 0h. The clusters contained genes that were further designated as either activated or suppressed by hypoxia (Fig. 7B, C).

We first looked at GSEA GO enrichments for all seven cases where a single GO term had a significant score for all three time points (Fig 7B). The predominant pattern was for gene activation, seen for 17 of the 21 data points. The only GO category that showed more suppression than activation was the *carbohydrate derivative biosynthetic process* (GO:1901137) from hmc7. Additionally, *circulatory system development* (GO:0072359) from hmc7 and *blood vessel development* (GO:0001568) from hmc3 were repressed at 1h, but then activated at 3h and 24h. All the others, including *cellular response to stress* (GO:0033554) from hmc9 and *cellular response to hypoxia* (GO:0071456) from hmc9 were activated at all three timepoints. Among the pathways identified, stress, hypoxia, and developmental signaling GO terms showed consistent activation.

We also looked at the most significant GO categories that were detected in only two of the three time points, displaying the top results for each pair group (Fig. 7C) or listing the full set (Supplementary Zip File 3, 4, 5). Notable GO terms detected in these categories include biologically and hypoxia-relevant terms such as response to oxidative stress, DNA repair, regulation of angiogenesis, and ncRNA processing. Other categories, such as organonitrogen compound biosynthesis, nucleoside triphosphate metabolism, and protein localization to organelle, have less obvious but potentially important metabolic roles. Additionally, several categories reflect cellular or organism developmental processes, such as neuron development, positive regulation of apoptosis, cell proliferation, and multicellular organism development. In terms of the direction of response, 3h and 24h maintained the same direction for all eight of the shared GO terms, while 1h and 24h showed different directions for two of the three shared GO terms. For 1h and 3h, two of the three GO terms maintained the same direction. These enrichments likely reflect the regulatory diversity of hypoxia-induced TF clusters and their involvement in a range of biological processes. Furthermore, these cluster groups helped to uncover subsets of TFs that may delineate regulatory modules for genomic responses to hypoxia.

### Clustering Reveals HIF1A and non-HIF1A Transcriptional Programs

Given the central role of HIF1A in modulating gene regulatory responses to hypoxia, we further analyzed our clusters with regard to HIF1A occupancy using both ChIP-seq and our footprinting data (Figure 8). To obtain a high confidence list of regions with the highest likelihood of having HIF1A occupancy, we used only the diff-MOA peaks that intersected with publicly available ReMap HIF1A ChIP-seq datasets. Having done this intersection for the diff-MOA GAIN and LOSS peaks separately (Fig. 8A, we observed a total of 439 GAIN peaks (18% of all GAIN peaks) overlapped HIF1A ChIP peaks, compared to 197 LOSS peaks (7% of all LOSS peaks). These ChIP-intersecting diff-MOA peaks were then mapped onto the hmc clusters (Fig. 8B; Supplementary Table S12). Nearly all of the GAIN peaks that intersected the HIF1A ChIP peaks (438 of 439, Fig. 8B) were limited to five clusters, hmc2,3,4,5, and 9. Similarly, all 197 of the LOSS peaks that intersected the HIF1A ChIP peaks were limited to the other five clusters, hmc1,6,7,8, and 10. To obtain an even higher confidence subset of HIF1A-related peaks, we further filtered the peak lists to obtain motifs (Supplementary Zip File 6) that also had the DNA sequence of the HIF1A motif, vdACGTGh from HOCOMOCOv11 (Kulakovskiy et al., 2018). These triple-hit regions (diff-MOA, HIF1A ChIP, and HIF1A motif; Fig. 8B, grey in bar graphs), were over-represented in the GAIN group which had a z-score of 4.19, but not in the LOSS group which had a (z-score = –1.07) (Supplemental Table S12).

**Figure 8.**
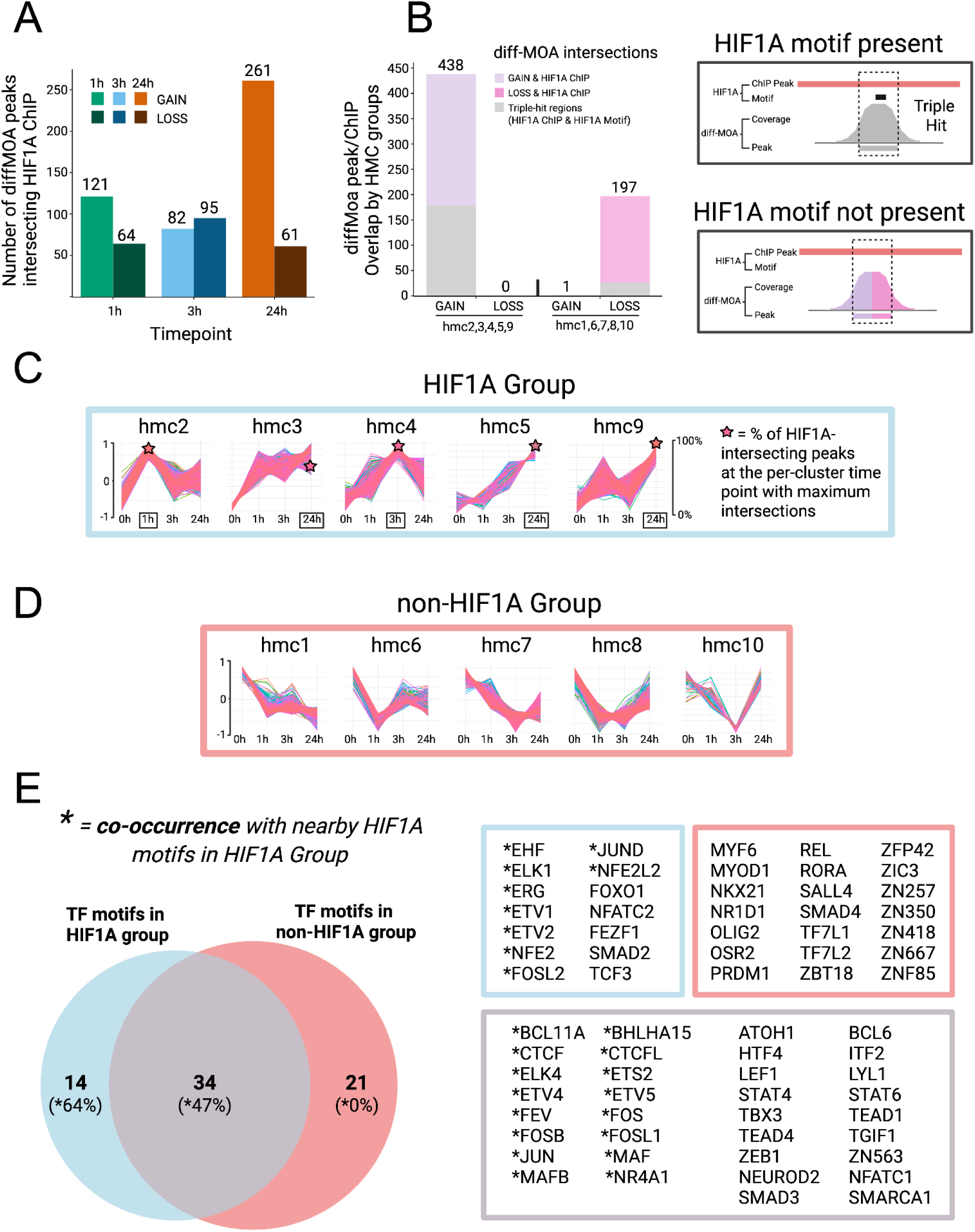
Classification into HIF1A and non-HIF1A groups, and motif co-occurrence analysis. MOA clusters were analyzed and grouped with regard to HIF1A occupancy using intersections of diff-MOA peaks with HIF1A ChIP-seq and HIF1A motif data. **(A)** Chart of the number of diff-MOA peaks that intersect with ReMap HIF1A ChIP-seq peaks, sorted by time and diff-MOA footprint type, GAIN or LOSS. **(B)** Chart showing the delineation of clusters (listed below the graph) into two groups according to diff-MOA footprint type, gain or loss. Each bar is further subdivided to show the proportion of diff-MOA peaks that intersect only HIF1A ChIP-seq peaks (lavender for GAIN, pink for LOSS) and those that also have the HIF1A motif (grey). The schematics depict the three-feature regions where the diff-MOA peak overlaps both HIF1A ChIP and HIF1A motif (grey) or the two-feature regions where the diff-MOA peak only intersects the HIF1A ChIP peak but not the motif (lavender for GAIN, pink for LOSS). **(C)** Classification of "hmc" clusters into HIF1A and non-HIF1A groups based on ChIP validation and HIF1A motif intersection greater than expected by chance (z-score > 2). The HIF1A group is shown with timepoints labeled on the x-axis for each cluster, left y-axis (as in Fig. 7A), and the right y-axis represents the percentage of HIF1A-intersecting peaks in a cluster. The star marks the percentage and the timepoint that makes up the greatest proportion of intersections (boxed on x-axis) with HIF1A ChIP for a given cluster. **(D)** The non-HIF1A group of clusters, which show a general downward trend for occupancy relative to 0h. **(E)** Other non-HIF1A TF motifs found in clusters from the two groups. The Venn diagram shows 21 non-HIF1A-group TFs (found in at least two clusters), 14 HIF1A-group TFs (found in at least three clusters), and 34 shared TFs. The unique and shared TF motifs are listed in the boxes, along with asterisks to indicate those which co-occur (+/- 50 bp) with the HIF1A motif. Percentages in the Venn diagram represent the proportion of TF motifs in each group that co-occur with the HIF1A motif.

This analysis allowed us to subdivide the clusters into two groups, the HIF1A Group (Fig. 8C) and the non-HIF1A Group (Fig. 8D), distinguished by intersection with HIF1A motifs and validated by correspondence with HIF1A ChIP peaks. These classifications align with the general expectation for HIF1A involvement in that the HIF1A Group exhibited common tendencies to show increased MOA signals over time relative to normoxia. Within the HIF1A Group, we found that the highest time point for MOA coverage showed strong concordance with the highest frequency of HIF1A ChIP peak intersection (indicated by stars in Fig. 8C). For instance, in hmc5 and hmc9, nearly all of the gain-of-footprint peaks that intersect HIF1A ChIP are found at the 24h timepoint, which corresponds to the maximum footprint timepoint in that cluster. Similarly, the footprint cluster maxima from hmc2 and hmc4 are found at 1h and 3h, respectively, matching their maximum footprints (Fig. 8C, Supplementary Table S12). In hmc3, the maximum HIF1A ChIP peaks are split between 3h (24%) and 24h (68%), which roughly mirrors the gain-of-footprint MOA-seq normalized read coverage maxima. Conversely, the non-HIF1A Group exhibited the opposite tendencies, showing reduced MOA signals resulting from hypoxia.

We also wanted to examine published endothelial PAS Domain Protein 1/Hypoxia-Inducible Transcription Factor 2A (EPAS1/HIF2A) ChIP-seq data (Fig. S7; Supplementary Table S12) but two important caveats precluded this analysis. First, the EPAS1/HIF2A motif (EPAS1/HIF2A: vdACGTGhh) is highly similar to that of HIF1A (HIF1A: vdACGTGh). Secondly, the vast majority (82%) of the EPAS1/HIF2A ChIP-seq peaks overlap with HIF1A ChIP-seq peaks (Fig. S7A).

To further investigate transcription factor motif composition across the two cluster groups defined by HIF1A association, we analyzed motif enrichment using the TF motifs (Fig. 7A) from the cluster-specific analysis. Motifs were listed if they appeared in at least three clusters from the HIF1A group (hmc2, 3, 4, 5, and 9) or at least two clusters from the non-HIF1A group (hmc1, 6, 7, 8, and 10) (Supplementary Table S13). We found 14 motifs unique to the HIF1A group, 21 unique to the non-HIF1A group, and 34 shared between both groups (Fig. 8E). Representative motifs unique to the HIF1A group include those for Nuclear Factor, Erythroid 2 (NFE2) and NFE2 Like BZIP Transcription Factor 2 (NFE2L2), both of which are known to be associated with hypoxia responses. In contrast, TF motifs unique to the non-HIF1A group include several motifs that as a group are substantially less-well linked to hypoxia.

Given that HIF1A does not act alone, we wanted to find other TF motifs that could be defined by their co-occurrence frequencies relative to vdACGTGh HIF1A motifs (Kulakovskiy et al., 2018) For this analysis, we defined co-occurence as motifs located within ±50 bp of a HIF1A motif and located within a diff-MOA peak (Supplementary Zip File 7). This analysis was performed separately for the HIF1A and non-HIF1A groups. In the HIF1A group, 64% of the TF motifs were unique to this group (e.g., ETS Transcription Factor ELK1 (ELK1), ETS Variant Transcription Factor 1 (ETV1)) and 47% of them were shared with the non-HIF1A group (e.g., MAF BZIP Transcription Factor B (MAFB), ETS Proto-Oncogene 2, Transcription Factor (ETS2)) (E-value < 0.05). In contrast, none of the motifs under diff-MOA peaks from the non-HIF1A group were significantly found within ±50 bp of HIF1A motifs, as expected from our classification scheme.

In summary, we demonstrate the use of MOA-seq with MNase footprinting to study small particle dynamics in HUVECs under hypoxic stress, generating a time-resolved atlas of over 5,000 genomic regions exhibiting significant cis-regulatory changes. This approach enabled high-resolution mapping of transcription factor occupancy and revealed widespread chromatin remodeling across the hypoxic time course. By clustering these dynamic footprints, we identified TFs or groups of TFs that may reflect distinct and multiple regulatory modules. Integration with HIF1A ChIP and motif analyses further partitioned these clusters into two larger groups, HIF1A and non-HIF1A, both of which likely contribute to the broader transcriptional architecture of the hypoxic response.

## Discussion

Deciphering the regulatory logic of gene expression requires not only identifying accessible chromatin regions, open chromatin, or nuclease hypersensitive sites but also pinpointing the precise sites of TF occupancy within those regions. Although genome-wide assays such as DNase-seq, ATAC-seq, and traditional MNase-based methods have mapped accessible chromatin landscapes with broad resolution, they often lack the precision needed to resolve individual TF binding events, particularly within densely occupied *cis*-regulatory domains (Baek & Sung, 2016; Vierstra et al., 2020). Similarly, ChIP-seq—while widely used—remains limited by antibody availability and specificity with peaks that typically span 100 bp or more (Luo et al., 2022).

As reported here, our MNase-based MOA-seq addresses this challenge by enabling high-resolution footprinting of TF occupancy across the genome, independent of antibody-based enrichment (Fig. 1). Previous applications in plant systems demonstrated that analysis of small fragments from light chromatin digests could detect thousands of discrete footprints (Savadel et al., 2021), enabling genetic analysis of cistrome dynamics (Engelhorn et al., 2025). In this study, we extend the utility of

MOA-seq to the human genome, demonstrating that it retains this high-resolution capacity, as similarly described in mammalian systems (Carone, 2025; Esnault et al., 2021). In normoxic HUVECs, MOA-seq footprints were sharply localized to transcription start and end sites and aligned closely with ENCODE candidate cis-regulatory elements (cCREs), validating its specificity for active regulatory regions (Fig. 2A–C). Strikingly, over one-third of high-confidence peaks fell outside of annotated cCREs or DNase I hypersensitive sites, revealing a previously underappreciated layer of regulatory potential (Fig. 2F–G). Furthermore, MOA-seq signal peaks were tightly centered on known transcription factor binding motifs and showed greater positional concordance with the cognate motifs than the actual ChIP-seq summits, highlighting its value for resolving the fine-scale mapping of TF occupancy (Fig. 3).

While MOA-seq provides valuable insights into transcription factor occupancy, several limitations or caveats should be noted. The particles captured may include non-transcription factor complexes, such as subnucleosomal structures or architectural proteins (Brahma & Henikoff, 2019; Ohno et al., 2019), and fragment size constraints limit the detection of larger complexes like the polycomb repressive complexes. The inability to uniquely map small fragment libraries may contribute to some false negatives. Furthermore, the redundancy of cis-regulatory motifs complicates precise transcription factor assignment, especially for heterodimeric or structurally similar factors such as families of TFs that bind the same sequences. As a pilot study, these results could benefit from deeper sequencing or more replicates, both of which could improve the statistical power to further validate or increase the total number of regions that pass quality control and statistical cutoffs used. Lastly, our findings are based on a single cell type, HUVECs, but showed good agreement with extensive TF and GO datasets from a large range of cell types and experimental systems.

In addressing the long-standing gap between accessible chromatin maps and high-resolution TF occupancy profiles, our study illustrates a robust methodology for capturing high-resolution genomic responses which can change over time as promoters are rapidly or slowly remodeled in response to hypoxia throughout the stress period (Suzuki et al., 2018). By cataloging cistrome occupancy at multiple timepoints, this single-assay approach helps establish a mechanistic framework for understanding how endothelial cells orchestrate complex, multi-phase responses to oxygen deprivation—insights not readily captured by single studies based primarily on transcriptomic or ChIP-based approaches.

To more precisely characterize these dynamic regulatory shifts, we examined how MOA-seq footprints change for the various sub-classes of previously defined candidate cis-regulatory elements, the ENCODE cCREs. Hypoxia-induced gain-of-footprint peaks were most prominent at CTCF-only and promoter-like sequence regions at 1 and 24 hours (Fig. 4D), suggesting that early hypoxia sensing and later-stage adaptation engage architectural proteins and transcriptional initiators in a biphasic manner (Kindrick & Mole, 2020). This pattern is consistent with a dual role for CTCF as an insulator and a scaffold for enhancer-promoter interactions shown to be required to activate hypoxia-specific gene networks (Kakani et al., 2025; Ren et al., 2017).

Conversely, loss-of-footprint peaks following hypoxia exposure were enriched at DNase I–H3K4me3-marked promoters across all time points (Fig. 4D), indicating early, transient, or sustained loss of gene activity (Batie et al., 2019). However, any singular MOA-seq footprint region alone cannot be definitively interpreted as corresponding to activators, repressors, or other DNA-binding factors. Thus, while some loss of footprint events may reflect reduced transcriptional activity, others, particularly those associated with upregulated genes, may also be consistent with loss of repressive factors.

Interestingly, many of dynamic sites we mapped lay outside ENCODE-annotated cCREs, yet were associated with genes enriched in pathways such as VEGFA–VEGFR2 signaling, G2–M checkpoint control, and TGF-β signaling (Figure 4D–E, 5A). These findings corroborate those from prior studies reporting the existence of novel Vascular Endothelial Growth Factor A (VEGFA)-responsive regulatory elements in endothelial cells (B. Zhang et al., 2013).

To functionally anchor these chromatin changes, we showed that genes with hypoxia-altered cistrome signatures were enriched in expected stress pathways, even without reference to transcriptomic data. Moreover, genes showing both occupancy shifts and differential expression were significantly more enriched for hypoxia-relevant pathways than those defined by either dataset alone (Fig. 5C), reinforcing the idea that chromatin accessibility dynamics often precede and help organize transcriptional outcomes (Batie et al., 2019; Liu et al., 2020). Motif analysis further supported this idea, with gain-of-footprint peaks being enriched for FOS, JUN, and CTCF motifs (Fig. 6A). Such observations are consistent with prior findings that stress-responsive factors such as AP1 help fine-tune hypoxia-induced transcriptional programs (Villar et al., 2012). Their presence at 1 and 24 hours, but not at 3 hours, suggests a biphasic response involving an early wave of TF engagement, a transient restructuring phase, and a second wave of TF engagement during adaptation. In contrast, loss-of-footprint peaks lacked strong enrichment for these major motif families. The exception was at 3 hours, where loss-of-footprint peaks were enriched for CTCF motifs, possibly reflecting hypoxia-induced CTCF-vacated regions (Fig. 6B).

The nature of this approach yields vast datasets of binding sites, implicating similarly large numbers of DNA-binding transcription factors. Although we have shown that global chromatin changes are detectable at specific regions enriched for cis-regulatory elements, the extensive amount of variation in gains and losses of site-specific footprint regions creates challenges for understanding how these genomic processes might be coordinated. Prior work has highlighted that hypoxia-responsive gene regulation is influenced by both the pre-existing epigenetic landscape and dynamic chromatin remodeling, which together modulate and extend the transcriptional response (Batie & Rocha, 2020; Kindrick & Mole, 2020). Moreover, early changes in histone modifications, such as H3K4me3 and H3K36me3, have been shown to predict downstream hypoxic transcriptional responses (Batie et al., 2019). However, it remains unclear how regulatory elements are integrated into temporally structured transcriptional programs, or how HIF1A coordinates with other factors to execute these responses (Villar et al., 2012; Yfantis et al., 2023). To address these questions and systematize the MOA-seq datasets, we decided to take a clustering approach, subdividing the thousands of peak regions into biologically informative co-varying groups, each enriched for smaller numbers of TF motifs (Fig. 7A).

The clustering of MOA-seq peak signal patterns over time yielded ten hypoxia MOA clusters (Fig. 7A). Among the clusters, hmc2, 3, 4, 5, and 9 showed increased binding relative to normoxia (Fig. 8B), while hmc1, 6, 7, 8, and 10 exhibited decreased binding (Fig. 8D). This stratification is consistent with the expected role of HIFA as a master regulator of the hypoxic response. Under low oxygen conditions, HIF1A becomes stabilized and translocates to the nucleus, where it drives widespread transcriptional activation by binding hypoxia response elements, the HREs (Kaluz et al., 2008).

Accordingly, clusters that showed trends towards gaining footprints after hypoxia showed strong enrichment for HIF1A motifs within ChIP-bound regions (Fig. 8B), supporting the interpretation that these modules are direct targets of HIF1A-mediated activation. In contrast, clusters tending to show decreased or loss of footprints after hypoxia did not exhibit significant enrichment for HIF1A motifs within ChIP-bound regions, even though some such sites were present. The presence of a few HIF1A motifs in these loss-of-footprint clusters may reflect the high genomic prevalence of HREs, numbering over one million genome-wide, but with less than 1,000 identified as HIF1A-bound from ChIP-seq experiments (Mole et al., 2009; Schödel et al., 2011).

Notably, due to the limited number of EPAS1/HIF2A ChIP-seq peaks available from consolidated external datasets and their extensive overlap with HIF1A ChIP peaks, we were unable to reliably distinguish EPAS1/HIF2A-specific binding from that of HIF1A in our analysis (Fig. S7). This reflects a broader challenge reported in the literature, where the ability to separate HIF1A and EPAS1/HIF2A binding events is limited by their shared HRE motif preference and substantial *in vivo* overlap, particularly in endothelial cells (Mole et al., 2009; Schödel et al., 2011; Smythies et al., 2019). While isoform-specific binding does occur (Bartoszewski et al., 2019), many target regions appear to be co-occupied, making it difficult to definitively assign regulatory activity to one isoform over the other.

By leveraging the quantitative variation in gain-of-footprint peaks at different timepoints within a cluster, we could see a clear concordance between the timepoint with the most HIF1A ChIP peaks and the highest level of MOA-seq signals for that cluster (Fig. 8C). For example, 88% of the HIF1A ChIP sites in hmc2 are found at 1h. Similarly, 95% of the HIF1A ChIP sites in hmc4 are found at 3h, and 99% of the HIF1A ChIP sites in hmc9 are found at 24h. These trends (denoted by stars in Fig. 8C) strongly suggests that HIF1A binding is a major contributor to the clustering patterns detected by our MOA-seq data in the five clusters we defined as the HIF1A group.

The concordance of HIF1A and cistrome occupancy clusters also serves to inform our understanding of the cluster-specific biological pathways. For instance, the transcriptional programs in hmc2 and hmc4 reflect coordinated early cellular reprogramming in response to hypoxia (Supplementary Zip File 3 and 4). Both 1 hour and 3 hour time points were marked by strong upregulation of pathways involved in catabolic remodeling, protein turnover, biosynthesis, and growth-related signaling—highlighting the cell’s immediate need to adapt metabolically and transcriptionally to low oxygen stress. This early phase supports proteostatic control and energy redistribution, laying the groundwork for survival under hypoxic conditions (Weckmann et al., 2018). In particular, the 3 hour response captured in hmc4 showed enhanced activity in biosynthesis, proteolysis, and supporting metabolic pathways—mechanisms that reinforce endothelial cell survival by supplying energy and biosynthetic precursors during early hypoxia (Liang et al., 2009). Alongside this broad activation, suppression of DNA repair pathways (Fig. 7C) coincided with the emergence of a mixed pro-proliferative and apoptotic transcriptional state, which has been shown to represent an intermediate phenotype at 3h of hypoxia that precedes a transition to anti-proliferation signaling during prolonged hypoxia in HUVECs (Liang et al., 2009).

As hypoxia progresses, endothelial cells undergo a phenotypic shift marked by repression of cell cycle progression and increased activation of apoptotic and stress-response pathways. Notably, our hmc9 cluster is dominated by maximal footprint signals at 24 hour, likely exemplifying this chronic hypoxia phase. Gene set enrichment in these clusters revealed sustained activation of hypoxia signaling, apoptosis, and suppression of cell cycle progression at the G1/S checkpoint—hallmarks of long-term cellular reorganization under oxygen deprivation (Supplementary Zip File 5) (Bartoszewski et al., 2019; Lee et al., 2020; Scheurer et al., 2004). These findings mirror previous observations that the early pro-growth and apoptotic transcriptional state seen around 3 hours transitions into a more definitive growth-suppressive phenotype by 8 hours and beyond (Liang et al., 2009). This is further supported by the broader overlap of pathways between 3 and 24 hours—including stronger enrichment for cellular response to hypoxia, consistent upregulation of vascular remodeling, which was suppressed at 1 hour (Fig. 7B), and angiogenesis, which was not present at the earliest time point (Fig. 7C). These results together with our cistrome response data support a biphasic regulatory model characterized by an initial wave of transcription factor engagement, a transient regulatory checkpoint at 3 hours, and a subsequent reactivation during prolonged hypoxia, as inferred from the cCREs with shifting distribution of gain and loss peaks over time. These results serve as reminders of the importance of temporarily resolved genomic studies, such as the initial minutes or hours following hypoxia, which may be key to elucidating multiple and overlapping genome response pathways.

Given that HIF1A does not act alone (Villar et al., 2012), a major challenge remains to fully identify and validate co-activators that collectively comprise the repertoire of HIF1A-mediated responses. Although chromatin accessibility plays a key role in guiding HIF binding by favoring regions with basal transcriptional activity and DNase I hypersensitivity, this alone does not fully explain HIF target selectivity, as many accessible loci with HRE motifs remain unbound or uninduced under hypoxia (Villar et al., 2012). Prior studies have shown that transcription factors such as AP-1 can co-bind near HIF sites and modulate hypoxic gene induction, with mutational analyses confirming their functional relevance in fine-tuning transcriptional output (Villar et al., 2012). These co-factors, many of which are also activated by hypoxia, likely contribute in a combinatorial and context-dependent fashion (Villar et al., 2012). In principle, our study should capture signatures of other DNA-binding proteins that interact nearby or directly with HIF1A. We observed co-occurrence of AP-1 family transcription factor motifs, including MAFB and FOS Like 2, AP-1 Transcription Factor Subunit (FOSL2), in proximity to HIF1A motifs within biologically relevant clusters of our HIF1A group (Fig. 8D). Specifically, hmc2 and hmc9, which we associate with acute and chronic hypoxic responses respectively, exhibit GSEA signatures that transition from early proliferative signals to apoptosis under prolonged hypoxia. This pattern is consistent with the modulatory influence of AP-1 across the hypoxic time course (Yadav et al., 2017).

Additional notable transcription factor motifs co-occurring with HIF1A include those of the ETS family, such as ELK1 and ETV4, which enhance HIF-mediated transcription of genes involved in angiogenesis and cellular adaptation (Ohnishi et al., 2013; Wollenick et al., 2012), and NFE2L2/NRF2 and MAF BZIP Transcription Factor (MAF) family members, which converge with HIF1A at oxidative stress-responsive elements to integrate redox and hypoxic signaling in a context-specific manner (Bae et al., 2024; Zheng et al., 2023). Within our hypoxia-associated cluster group, we also observed several transcription factor motifs implicated in the hypoxic response that did not co-occur spatially with HIF1A motifs (Fig. 8D; TFs without asterisks in the blue or grey boxes), yet exhibited temporal patterns of occupancy that paralleled HIF1A binding patterns. One such factor is Forkhead Box O1 (FOXO1), which has been shown to promote antioxidant gene expression through AMP-activated protein kinase (AMPK)-dependent nuclear translocation under hypoxia (Awad et al., 2014). Although FOXO1 motifs were not enriched near HIF1A-bound regions, their presence in clusters with coordinated activation suggests they participate in parallel and temporally synchronized regulatory circuits, potentially driven by shared upstream signals such as redox or energy stress. These programs may complement HIF1A-mediated transcription without direct co-occupancy, but exhibit features of co-activation or genetic interaction impinging on the same phenotypic pathways. Importantly, our inability to definitively distinguish HIF1A from EPAS1/HIF2A occupancy at shared hypoxia response elements raises the possibility that some motif enrichments—particularly at later time points such as 24 hours—may reflect partners or targets of EPAS1/HIF2A rather than HIF1A.

In contrast to the HIF1A group, the non-HIF1A clusters (hmc1,6 7,8,10) represent distinct classes of cistrome responses that exhibit reduced binding under hypoxic conditions relative to normoxia. Despite this overall loss of MOA-seq footprinting, these clusters still displayed five distinct temporal patterns. The transcription factor motifs enriched in these clusters included regulators such as Myogenic Differentiation 1 (MYOD1), Myogenic Factor 6 (MYF6), Oligodendrocyte Transcription Factor 2 (OLIG2), RAR Related Orphan Receptor A (RORA), and multiple zinc finger proteins (e.g., Zinc Finger And BTB Domain Containing 18 (ZBTB18), Zinc Finger Protein 257 (ZNF257), Zinc Finger Protein 667 (ZNF667)), many of which are involved in lineage specification, cell fate determination, and context-dependent gene repression (Casey et al., 2018; Schmitges et al., 2016). While these factors may generally relate to development, they may reflect a more general set of controls for turning off genes and pathways as part of stress adaptation responses.

Several of the transcription factors enriched in the non-HIF1A group, such as MYOD1 and OLIG2, have been previously linked to hypoxia signaling through indirect or cell-type–specific interactions with HIF1A (Allan et al., 2021; Cirillo et al., 2017). MYOD1 can influence HIF1A activity through MyoD-mediated activation of the noncanonical Wnt/β-catenin pathway during myogenesis, while OLIG2 and HIF1A cooperatively regulate gene expression in oligodendrocyte progenitor cells under hypoxic conditions, suggesting broader roles for these factors in modulating the hypoxic response outside of direct enhancer co-occupancy. Others, such as Zinc Finger E-Box Binding Homeobox 1 (ZEB1) and Transcription Factor 7 Like 1 (TCF7L1), may represent novel hypoxia-responsive regulators that function independently of HIF1A (Yoshimoto et al., 2019; B. Zhang et al., 2019). ZEB1 is induced by hypoxia and contributes to epithelial-to-mesenchymal transition, a process in which HIF1A has also been implicated, while TCF7L1—found enriched in cluster 6—promotes oxidative stress responses via the Kelch Like ECH Associated Protein 1 (KEAP1)/NRF2 pathway and may play a role in early transcriptional priming. Finally, 34 transcription factor motifs were shared between the HIF1A and non-HIF1A groups, including ETS2, MAFB, and SMAD3, suggesting that crosstalk or sequential coordination may exist between these regulatory programs. The presence of shared motifs implies that some transcription factors may participate in both HIF1A-bound and HIF1A-independent modules, potentially integrating canonical and noncanonical hypoxic signals depending on chromatin context, TF availability, or temporal stage of the hypoxic response.

Future studies should integrate MOA-seq with complementary epigenomic and transcriptomic datasets to build a more complete model of hypoxia-driven gene regulation. Expanding analyses to additional cell types, time points, or environmental conditions holds great promise for further advancing our knowledge of chromatin remodeling and transcription factor network dynamics across biological contexts.

Functional validation of novel regulatory elements identified here will also be essential to define their roles in gene expression and disease progression. Taken together, this study demonstrates how cistrome occupancy profiling offers a high-resolution view of dynamic regulatory element engagement under hypoxia. Cistrome occupancy clustering over multiple time points bifurcated our hypoxia responsive regions into two large groups reflecting HIF1A and non-HIF1A, each composed of 5 clusters with largely distinct TF motifs and biological pathways. By capturing genomic responses in the form of small particle dynamics, this approach expanded our understanding of hypoxia-responsive networks and provided a new framework for exploring chromatin-based transcriptional control in health and disease.

## Materials and Methods

### MOA-seq library preparation and sequencing

HUVECs (ATCC, Cat. No. PSC-100-010) were cultured in endothelial growth media under standard culture conditions (5% CO₂, 95% room air, 37°C) and maintained between passages 5–7. Approximately 900,000 cells were seeded into 100 mm culture dishes and typically reached 90–95% confluency within 24–48 hours. Cells were certified as mycoplasma-free by the supplier; STR genotyping was not performed. At 90–95% confluency, cells were subjected to hypoxia (2% O₂) for 1, 3, or 24 hours.

Following hypoxia treatment, cells were formaldehyde-fixed and stored at –80°C until processing. To isolate nuclei, fixed cells from a 10 cm dish were thawed and resuspended in 500 µL Nucleus Isolation Buffer (NIB: 0.3 M sucrose, 2 mM MgOAc, 2.1% Triton X-100, 10 mM HEPES pH 7.4, 1 mM CaCl₂) and incubated at room temperature for 5 minutes. Cells were pelleted (5 minutes, 1000 × g), washed in fresh NIB, and resuspended in 500 µL NIB. A small aliquot (10 µL) was retained for microscopy checks of nuclear integrity. The remaining nuclei suspension was aliquoted into eight 1.5 mL tubes (60 µL per tube). Micrococcal nuclease (MNase) digestion was performed using a serial dilution series of MNase (starting at 1280 U/mL and decreasing two-fold to 20 U/mL). Each enzyme dilution (6.7 µL) was added to corresponding nuclei aliquots, and digestion proceeded at 37°C for 15 minutes. Reactions were stopped by adding 5 µL of 0.5 M EGTA-NaOH (pH 8.0). Decrosslinking was performed overnight at 65°C with 360 µL ddH₂O, 50 µL 10% SDS, 50 µL 1.5 M NaCl, and 5 µL 20 mg/mL Proteinase K. DNA was extracted through sequential phase separation with

Tris-saturated phenol, followed by phenol:chloroform (1:1), and chloroform alone. DNA was precipitated by adding 0.05 volumes of 3 M sodium acetate (pH 5.2), 2 µL 25 mg/mL linear polyacrylamide, and 2 volumes of –20°C ethanol. The mixture was incubated at –20°C for at least 30 minutes and then pelleted by centrifugation (20,000 × g, 15 minutes, 4°C). The pellet was washed in 70% ethanol, air-dried, and resuspended in 100 µL TE buffer containing 40 µg/mL RNase A. RNA was removed by incubation at room temperature for 1 hour, and DNA was re-precipitated as previously described. Final DNA pellets were resuspended in 30 µL TE buffer, quantified by spectrophotometry, and analyzed on a 1% agarose gel. DNA quality was further assessed using an Agilent 2100 Bioanalyzer, Q-PCR, and a Qubit 2.0 Fluorometer. Libraries were prepared using NEBNext® Ultra™ II DNA Library Prep Kits (Cat. E7103S). The indexed libraries were size-selected to enrich for sub-nucleosomal (<150 bp) genomic insert sequences and submitted for paired-end 150 bp reads on the Illumina HiSeq (FSU College of Medicine, Translational Science Lab). Two biological replicates were analyzed per condition.

For control MOA libraries, DNA was isolated from A549 human cells (ATCC, CCL-185), cultured in F12K media supplemented with fetal bovine serum, 1% L-glutamine, and 1% penicillin-streptomycin. DNA from A549 cells was extracted with Invitrogen™ TRIzol™ Reagent (as described in Chomczynski, 1993)(Chomczynski, 1993). Briefly, cells were harvested using trypsin and centrifuged at 500 × g for 5 minutes at 4°C to pellet cells.

The cell pellet was resuspended in TRIzol™ Reagent. After the addition of chloroform, samples were centrifuged at 12,000 × g at 4°C for 15 minutes. The aqueous phase, containing RNA, was discarded, and the DNA was precipitated from the interphase/organic layer with ethanol. The precipitated DNA was washed twice with 0.1 M sodium citrate in 10% ethanol and pelleted at 2000 × g at 4°C for 5 minutes. The pellet was washed in 75% ethanol and pelleted again at 2000 × g at 4°C for 5 minutes. The cleaned DNA pellet was resuspended in an elution buffer (10 mM Tris-HCl, 0.8 mM EDTA, pH 8.5). DNA quality was assessed via agarose gel electrophoresis, and concentrations were determined using a NanoDrop UV spectrophotometer. DNA yields were sufficient for multiple MNase digestions. MNase digestion conditions were optimized based on MOA-seq chromatin digestion, scaled to 1/64th concentration.

Digestion was performed to achieve a partial digest pattern similar to MOA-seq. Digest volumes were scaled to 60 µL to yield ∼1 µg of DNA per digest for Illumina library preparation. Digestion patterns were assessed via gel electrophoresis, and the light digest levels ideal for MOA-seq (as per Savadel et al., 2021) were selected as the heaviest digest levels that give a pattern of a nucleosomal ladder spanning the entire DNA fragment size range from monosomal bands (∼ 150 bp) to just below the size of undigested DNA fragments. Select digests matching these criteria are indicated in Figure 1 in the yellow asterisk-marked gel lanes. The selected samples were pooled to increase DNA input to 1 µg per sample. Library preparation for control MOA libraries followed the NEBNext Ultra II DNA Library Prep Kit for Illumina (NEB #E7103). After adapters were added, samples were flash-frozen, and library-specific index sequences were incorporated using the dual indices method by the NGS Library Facility at FSU. DNA libraries were sequenced with a target read depth of 80 million reads per sample and stored at –80°C until sequencing.

### MOA-seq data processing and fragment center coverage

The sequence data reads (in FASTQ format) were trimmed for adapters using Cutadapt (Martin, 2011) with default parameters. Because the sequence read chemistry generates overlapping reads, the trimmed paired-end reads for each biological replicate were subsequently merged using FLASh (Magoč & Salzberg, 2011) with the parameters “-t 40 --cap-mismatch-quals -M 150 -d -q”. The resulting merged read files were aligned to the human reference genome (hg38) using the bwa-mem2 mem function (Vasimuddin et al., 2019) with the parameters “-M -t 40.”

Alignment files (SAM) were converted to BAM format using SAMtools (H. Li et al., 2009), with additional filtering applied to retain only reads with a mapping quality score (MAPQ) of 20 or greater. PCR duplicates were also removed. The average fragment length was calculated using the stat function in SAMtools (H. Li et al., 2009), while the effective genome size was determined using unique-kmers.py (Crusoe et al., 2015) with the average fragment length and the hg38 genome FASTA as inputs. BAM files were converted to BED files using the bamToBed function in Bedtools (Quinlan & Hall, 2010). The alignment and pre-processing metrics are provided in Supplementary Table S1.

To achieve greater resolution, fragment center tracks (also referred to as ’frenters’ as described by Savadel et al., 2021) were created. In this process, each mapped read was shortened to its central 20 bp using a custom awk command. For reads with an uneven number of bases, the central base was taken and extended by 10 bp. For reads with an even number of base pairs, the central base was randomly selected from one of the two central bases and then extended by 10 bp. Bedgraphs were generated using the genomeCoverageBed function in Bedtools (Quinlan & Hall, 2010) with a counts-per-million (CPM) scaling factor to normalize for sequencing depth. Finally, BigWig files for UCSC Genome Browser visualization were produced using the bedGraphToBigWig utility (Kent et al., 2010), a UCSC binary tool.

### Within sample and differential peak calling

Macs3 was used for peak calling (Y. Zhang et al., 2008). Fragment center BED files for each biological replicate pair per treatment were used as input to the callpeak function of MACS3 with the following parameters “-p .001 --nomodel --extsize 20 --min-length 20 --max-gap 20 --keep-dup all.” The resulting peak sets were analyzed using the Irreproducibility Discovery Rate (IDR) framework (Q. Li et al., 2011) to assess data consistency, identifying reproducible peaks between biological replicates within each sample, referred to as within-sample peaks. In accordance with ENCODE ChIP-seq guidelines (Landt et al., 2012), we further evaluated data quality by assessing pooled pseudo-replicate consistency and self-consistency for each individual replicate (Supplementary Table S2). To minimize artifacts, within-sample peaks were filtered using the intersect function in Bedtools to exclude blacklist regions (Amemiya et al., 2019) and MOA control regions, defined by chromatin-free total DNA light digests with MNase.

To identify hypoxia-induced binding sites (gained, lost, or shared), we used the bdgdiff function in MACS3, comparing each hypoxia time point (1h, 3h, 24h) to the normoxic sample. Combined replicate BED files for each sample were used as input for bdgdiff analysis, with the minimum peak length and maximum peak gap parameters set to 20. The optimal likelihood ratio (LHR) cutoff was determined using the elbow method (Supplementary Table S2).

### MOA-seq enrichment analyses

We gathered various ENCODE-published datasets for MOA-seq enrichment analysis. Bulk RNA-seq gene quantifications (“An Integrated Encyclopedia of DNA Elements in the Human Genome,” 2012) (ENCFF892OHT) were obtained for endothelial cells and used to demonstrate the correlation between MOA normoxic coverage and gene expression at the transcription start site (TSS) and the transcription end site (TES).

Candidate cis-regulatory elements (cCREs) and DNase I data from multiple cell types (Sabo et al., 2004, 2006) were collected for their genomic coordinates and served as key features for plotting local average normoxic MOA coverage and analyzing normoxic peak base pair intersections. To evaluate the base pair intersections of normoxic peaks with ENCODE cCREs and DNase I data, a probability distribution was generated by intersecting shuffled normoxic peak coordinates with the published dataset genomic coordinates using Bedtools 100 times. This approach determined the expected number of base pairs that would intersect by chance. The expected base pair observations were plotted to create a probability distribution, which was then used to calculate a z-score (Supplementary Table S3). The z-score quantified how many standard deviations the observed number of intersected base pairs deviated from the expected number. The genomic annotation of normoxic peaks was performed using the annotatePeak function in ChIPseeker (Q. Wang et al., 2022; Yu et al., 2015).

ChIP-seq peaks of various transcription factors for the HUVEC cell type were obtained from the ReMap 2022 ChIP-seq database (Hammal et al., 2022) (Experiments and peak counts listed in Supplementary Table S14). Their genomic coordinates were used as central features for plotting the local average normoxic MOA coverage. The peak-to-flank enrichment ratio was calculated by dividing the MOA-seq signal at the peak summit by the average signal across the 50 bp flanks on either side of the coverage window (totaling 100 bp). The canonical motif for the ERG, JUN, and ETS1 ReMap ChIP-seq peak sets was identified using the SEA tool in the MEME suite (Bailey et al., 2015) and used as the central feature for plotting local average normoxic MOA coverage. ChIP-seq FASTQ files for each of these transcription factors, derived from a single HUVEC experiment (S. Wang et al., 2019), were aligned to the hg38 reference genome using BWA-MEM2 with default parameters. The resulting ChIP-seq coverage was plotted alongside the normoxic MOA coverage, centered on the canonical motif for each respective TF. For this, the y-axis represents the average reads per million at each base pair, normalized to the minimum value for each profile. Coverage heatmaps were generated using Deeptools plotHeatmap (Ramírez et al., 2014), focusing only on ChIP-seq peaks with non-zero MOA coverage within the 400-bp window.

Hypoxia-induced binding sites or diff-MOA were visualized using heatmaps generated with Deeptools plotHeatmap, overlaying hypoxia and normoxia coverage. Gain sites were sorted by hypoxia coverage values, while loss sites were sorted by normoxia coverage values. The diff-MOA from all time points were intersected with the gene bodies of protein-coding genes from GENCODE version 45 (Frankish et al., 2023) using Bedtools intersect, with gene start and end coordinates extended by 200 base pairs to capture potential regulatory elements downstream of the TSS or TES. These genes were termed diff-MOA genes and were then grouped based on whether the intersecting diff-MOA peak was located within a cCRE, outside a cCRE, or if the gene contained peaks in both regions (Supplementary Table S5). Any identified gene sets were submitted to Enrichr for transcription, pathway, and gene ontology enrichment analysis. To assess the statistical significance of the transcriptional pathways for the diff-MOA gene lists, we generated randomized size-matched gene lists for Enrichr input to simulate a random genomic background for comparison (Supplementary Table S5). The gain, loss, and shared diff-MOA peaks were also intersected with ENCODE cCREs to determine the percentage of peaks overlapping each cCRE subtype (Supplementary Table S4). Base pair intersections were performed as well, and the density was calculated by dividing the total number of base pairs intersecting a given cCRE subtype by the total number of base pairs comprising that subtype, then multiplying by one million to report density per megabase (Supplementary Table S4).

Gain and loss diff-MOA peaks by hour were submitted separately and in combination to Enrichr for pathway analysis (Supplementary Table S6). We then examined the relationship between diff-MOA and differentially expressed genes (DEGs) by identifying DEGs that also had a gain or loss diff-MOA peak. These subsets of genes were submitted to Enrichr for pathway analysis using the MSigDB Hallmark and WikiPathways Human libraries. To assess whether the diff-MOA-identified subset of DEGs exhibited stronger statistical enrichment, we created three size-matched gene subsets, randomly sampled from the list of DEGs for each time point. Each subset was submitted to Enrichr, and the average statistical significance from the three inputs was calculated to determine whether a random, size-matched subset of DEGs performed comparably (Supplementary Table S7).

### Hierarchical clustering analysis

All diff-MOA peaks, including gains and losses at 1h, 3h, or 24h of hypoxia, were merged and windowed at 30 bp centered on the merged peaks. Normalized read-depth coverage counts for each time point were calculated for each window using the coverage function in Bedtools (Quinlan & Hall, 2010). These normalized counts were standardized using z-score normalization for each window or peak across the three hypoxia time points and used as input for clustering analysis. A hierarchical dendrogram was generated using the hclust function in R. The optimal number of clusters (k = 10) was determined using the elbow method, where the within-cluster sum of squares (WCSS) stabilized and plateaued (Supplementary Table S9). The resulting 10 clusters (hypoxia MOA clusters or "hmc"s) were visualized using ggplot2 (Wickham, n.d.), with each line in the cluster graphs representing a single differential peak and its MOA coverage across the four sample time points. The unstandardized MOA coverage values for each hour and the genomic coordinates for each merged diff-MOA peak, organized by cluster, are provided in Supplementary Table S10.

### Motif discovery and analysis

Motif discovery, enrichment, and subsequent analyses utilized XSTREME from the MEME suite (Bailey et al., 2015). Default settings were applied to the diff-MOA peaks, with the Markov order set to M = 0 (Supplementary Zip File 1, Supplementary Table S8). Genomic sequences from the diff-MOA peaks were extracted using the Bedtools getfasta function and served as input for XSTREME. Motif enrichment analysis employed the HOCOMOCOv11 transcription factor motif database (Kulakovskiy et al., 2018). The top three motifs for each diff-MOA set, ranked by E-value, were then intersected with the corresponding ReMap ChIP-seq data, where available. Additionally, genomic sequences from each hmc cluster were processed by XSTREME to identify cluster-specific TF motifs (Supplementary Table S11, Supplementary Zip File 2). The motif enrichment tool SEA from the MEME suite identified HIF1A and EPAS1/HIF2A motifs in hypoxia-induced footprint gain and loss sites for 1h, 3h, and 24h (Supplementary Zip File 6). These sites were categorized based on their intersection with HIF1A and EPAS1 ReMap ChIP peaks, respectively, and their distribution within the hypoxia MOA clusters (Supplementary Table S12). To determine whether diff-MOA peaks overlapping HIF1A ChIP peaks were significantly associated with the motif, we subsampled differential MOA peaks 100 times, recorded the motif frequency as our expected, and performed a z-score analysis to determine if the observed motifs associated with the ChIP peaks were more frequent than expected by chance in our diff-MOA peaks without ChIP validation (Supplementary Table S12).

To identify motifs co-occurring near HIF1A motifs in either the HIF1A group clusters (hmc2,3,4,5,9) or the non-HIF1A group clusters (hmc1,6,7,8,10), we extracted the central base pair genomic coordinates of peaks containing HIF1A motifs in these cluster groups and extended the window by 50 bp on either side, resulting in a 101-bp window. These coordinates underwent analysis using the motif enrichment tool SEA from the MEME suite (Supplementary Zip File 7). From the prior motif analysis on each individual cluster, we identified common TF motifs between the clusters that make up our HIF1A group and our non-HIF1A group. Among these common TFs, those that co-occurred with HIF1A motifs in the immediate region were marked with an asterisk (Supplementary Table S13).

### Gene Set Enrichment Analysis

Differential gene expression analysis was performed using DESeq2 (Love et al., 2014) to process raw read counts and identify differentially expressed genes (DEGs). Genes with insufficient read counts across replicates were filtered out. DESeq2 was used to normalize the data and calculate log2 fold changes (Log2FC) for each condition (1h, 3h, 24h) relative to the baseline (0h). Significant DEGs were identified for each time point using an adjusted p-value threshold of ≤ 0.05. A combined list of DEGs was generated, and Log2FC values were retained for each time point for downstream analysis.

From this combined list, only DEGs associated with a defined cluster were included in subsequent analyses, as not all DEGs corresponded to a cluster. The selected DEGs were organized into their respective clusters, and gene set enrichment analysis (GSEA) (Mootha et al., 2003; Subramanian et al., 2005) was performed for each time point using the DEGs within a given cluster and their corresponding Log2FC values. The gseGO function from the clusterProfiler R package was used to identify enriched biological processes (BP ontology). Enrichment parameters included a minimum gene set size of 3, a maximum gene set size of 800, and a p-value cutoff of 0.05. For each cluster and time point, normalized enrichment scores (NES), p-values, and gene ratios were calculated, where gene ratios represented the proportion of genes contributing to enrichment within each gene set. The outputs of the enrichment analysis, including cluster-specific enrichment results for each time point, were exported as CSV files for further interpretation (Supplementary Zip File 3, 4, 5).

MOA processing and custom analysis scripts are available from https://doi.org/10.5281/zenodo.16595154 and have been deposited in the GitHub repository (https://github.com/as5906/HypoxiaMOA-HUVEC).

### Data Access

All MOA-seq data discussed in this publication have been deposited at NCBI SRA under accession number PRJNA1217882. MOA bigwig and peak files are available on Gene Expression Omnibus (ID: GSE289863).

## Supporting information

Singh--Bass_Supplemental_TABLE_S1

Singh--Bass_Supplemental_TABLE_S5

Singh--Bass_Supplemental_TABLE_S10

Singh--Bass_Supplemental_TABLE_S6

Singh--Bass_Supplemental_TABLE_S7

Singh--Bass_Supplemental_TABLE_S2

Singh--Bass_Supplemental_TABLE_S9

Singh--Bass_Supplemental_TABLE_S3

Singh--Bass_Supplemental_TABLE_S12

Singh--Bass_Supplemental_TABLE_S13

Singh--Bass_Supplemental_TABLE_S8

Singh--Bass_Supplemental_TABLE_S11

Singh--Bass_Supplemental_TABLE_S4

Singh--Bass_Supplemental_TABLE_S14

Singh--Bass_Supplemental_FILE_S1

Singh--Bass_Supplemental_FILE_S2

Singh--Bass_Supplemental_FILE_S5

Singh--Bass_Supplemental_FILE_S4

Singh--Bass_Supplemental_FILE_S3

Singh--Bass_Supplemental_FILE_S6

Singh--Bass_Supplemental_FILE_S7

## Acknowledgements

This work was supported by an NIH grant to MNG and HWB (NIH R01 HL177087); an NSF Plant Genome Research Program grant to HWB (NSF IOS 1444532, 2025811); a Phi Beta Kappa MJ Hay Award and Biological Science Faculty Award for Undergraduate Scholarship to AS; and an American Heart Association grant to GTD (Award No. 830166).

## Supplementary Figures

**Figure S1.**
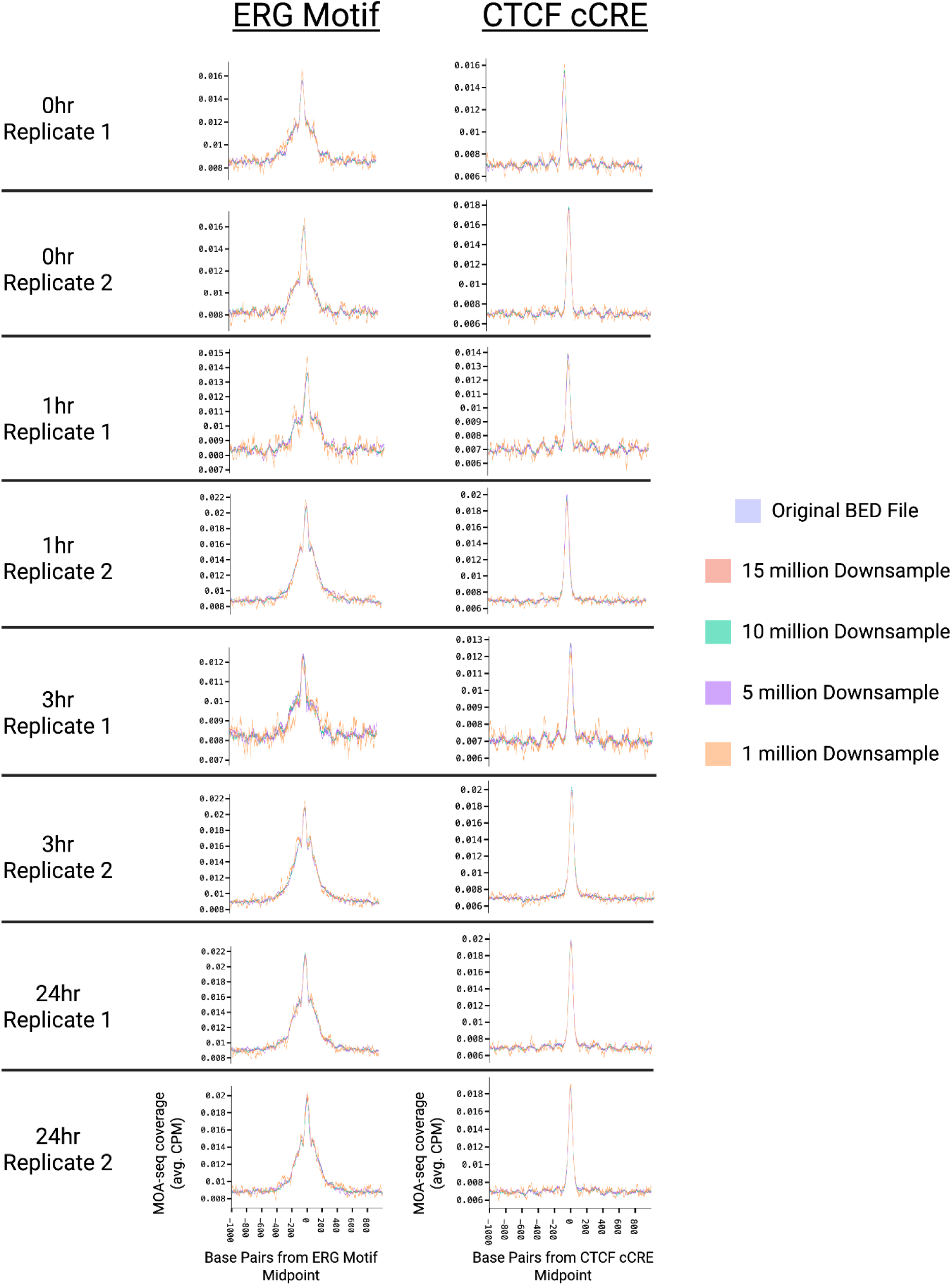
MOA-seq signal stability across read-depth downsampling. Aggregate MOA-seq coverage profiles (average counts per million, CPM) centered on ERG motif midpoints (left column) and CTCF candidate cis-regulatory elements (cCREs; right column) are shown for two biological replicates at 0, 1, 3, and 24 hours. For each panel, the original dataset (blue) is overlaid with profiles generated after random downsampling to 15 million, 10 million, 5 million, and 1 million reads (colored traces).

**Figure S2.**
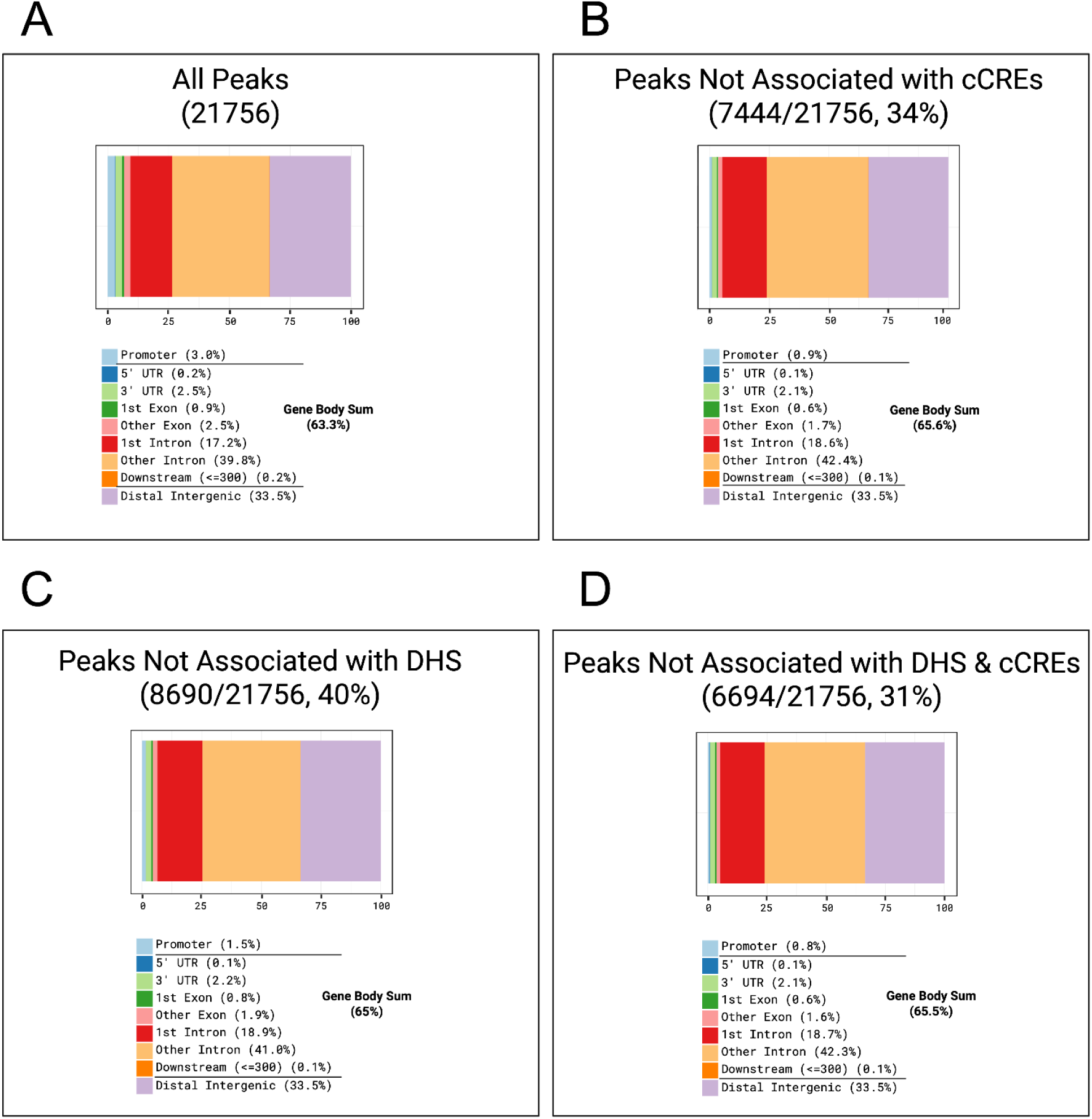
Genomic distribution of normoxic MOA peaks. Comparable genomic distribution of MOA peaks detected under normoxic conditions, annotated using ChIPseeker (Q. Wang et al., 2022; Yu et al., 2015). Peaks were classified by genomic context, including promoter (±200 bp from the TSS), 5′ UTR, 3′ UTR, exon, intron (first and other), downstream, and distal intergenic regions. **(A)** All normoxic peaks (n = 21,765, 100%). **(B)** Peaks not overlapping DHSs (n = 9,803, 45.0%). **(C)** Peaks not overlapping cCREs (n = 12,274, 56.4%). **(D)** Peaks overlapping neither DHSs nor cCREs (n = 7,202, 33.1%).

**Figure S3.**
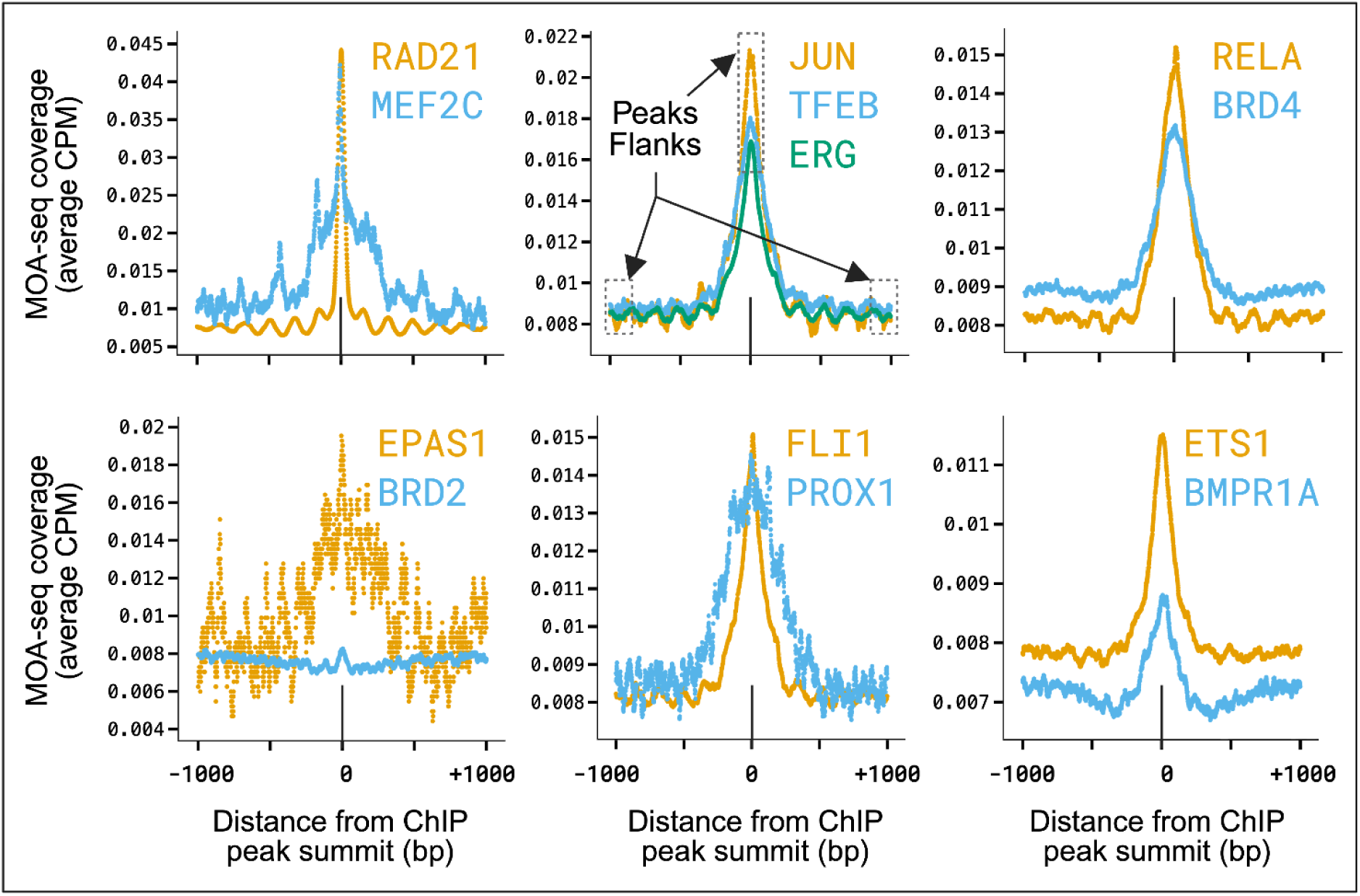
**MOA Coverage at ReMap ChIP summits from HUVEC experiments for 13 transcription factors**. Profile trend plots of local MOA frenter coverage for the individual TFs grouped primarily by signal height. Plots are centered on the ReMAP ChIP peak summit for each TF.

**Figure S4.**
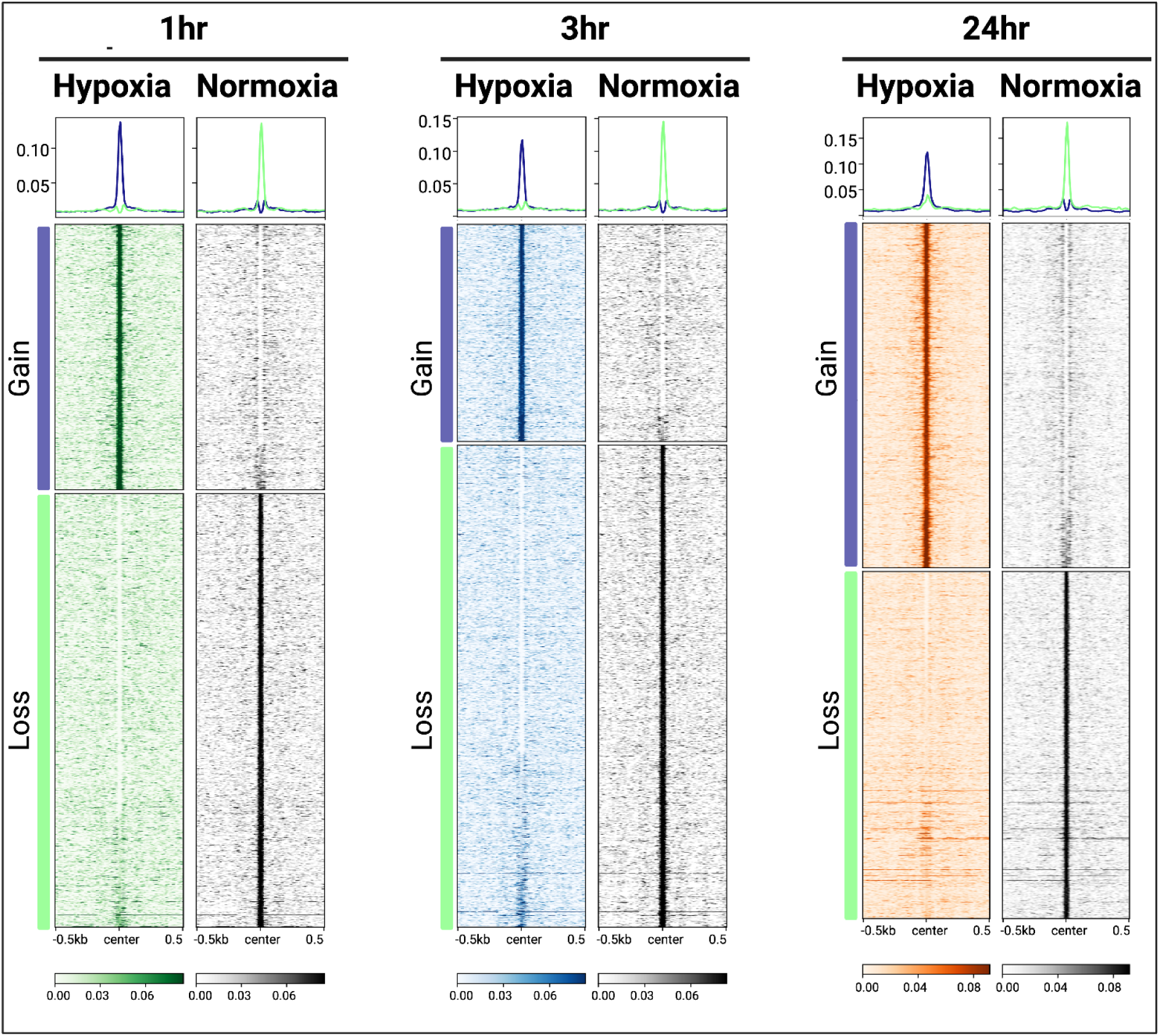
Heatmaps of MOA-seq frenter coverage at diff-MOA peaks across hypoxia time points. MOA-seq frenter coverage heatmaps of the diff-MOA (GAIN vs LOSS of footprints relative to normoxia) for each time point. Heatmaps were sorted by total MOA signal averaged over the 1 kb window centered on the peaks. In addition, sorting was done for diff-MOA gain sites using the hypoxic time point as the primary sort, and for diff-MOA loss sites, using the normoxic time point as the primary sort.

**Figure S5.**
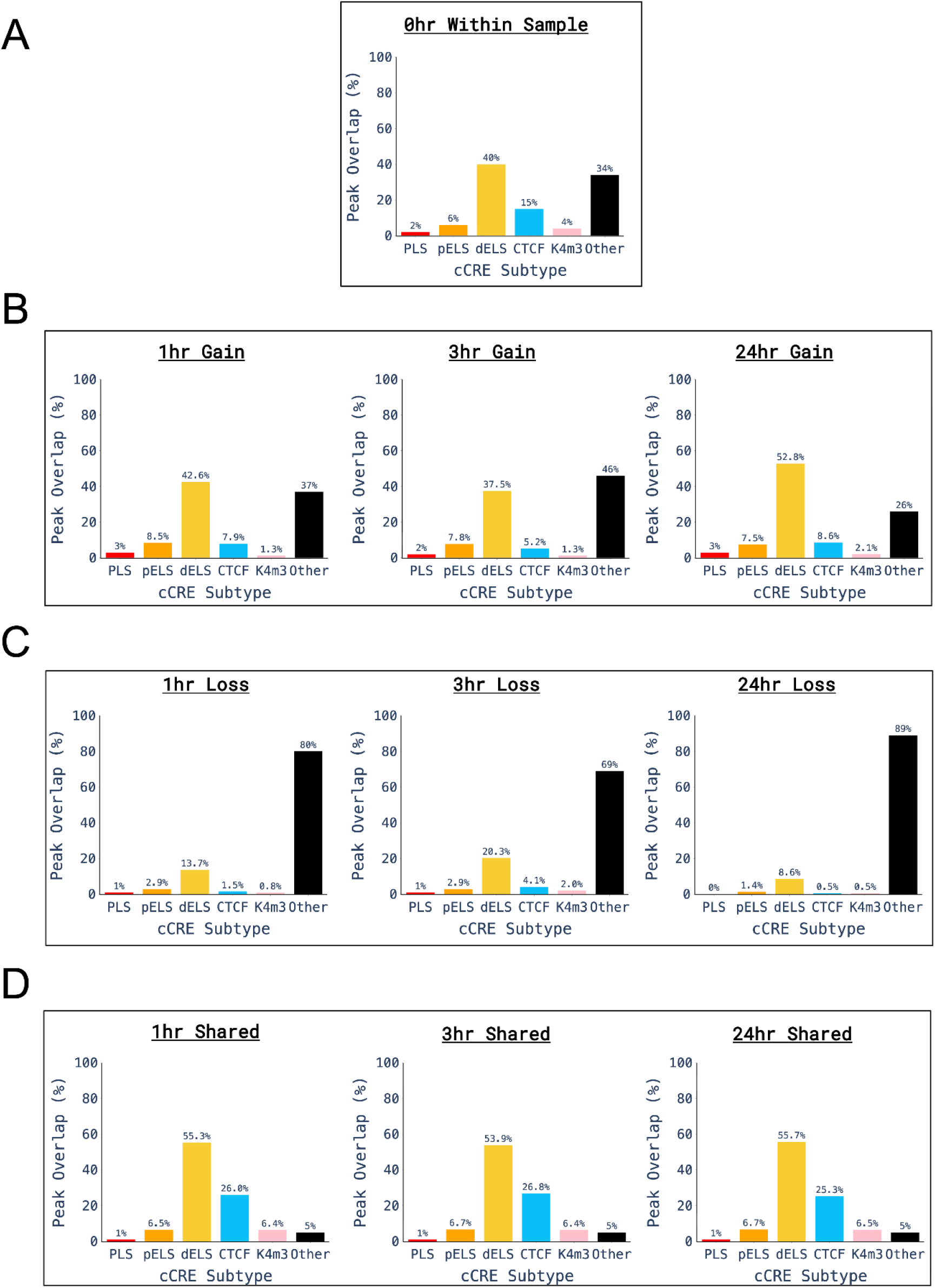
Distribution of diff-MOA peaks across ENCODE cCRE subclasses. Percentage of peaks overlapping ENCODE cCRE subclasses: PLS (promoter-like signature), pELS (proximal enhancer-like signature), dELS (distal enhancer-like signature), CTCF-binding sites, K4m3 (DNase I–H3K4me3), and Other. The “Other” category refers to peaks not overlapping any ENCODE cCRE subclass. **(A)** Normoxic MOA peaks. **(B)** diff-MOA gain peaks at 1h, 3h, and 24h. **(C)** diff-MOA loss peaks at 1 h, 3 h, and 24 h. **(D)** diff-MOA shared peaks at 1h, 3h, and 24h.

**Figure S6.**
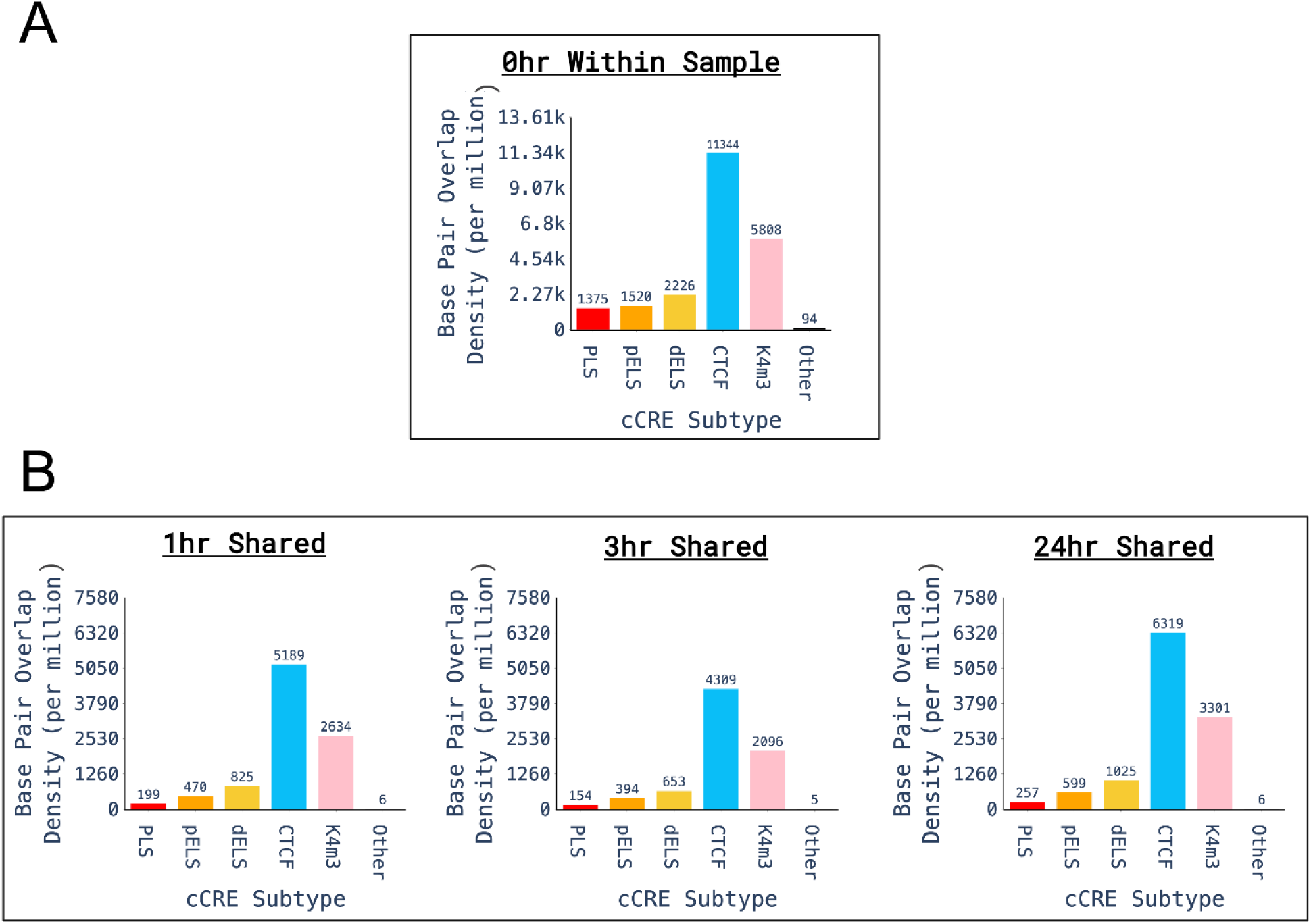
Base pair density overlap of normoxic and shared diff-MOA peaks across ENCODE cCRE subclasses. Base pair density overlap (per megabase) of MOA peaks across ENCODE cCRE subclasses: K4m3 (DNase I–H3K4me3), dELS (distal enhancer-like signature), pELS (proximal enhancer-like signature), and PLS (promoter-like signature), along with an additional category labeled “Other” representing peaks not overlapping any cCRE subclass. **(A)** Normoxic MOA peaks. **(B)** Shared diff-MOA peaks at 1h, 3h, 2 h.

**Figure S7.**
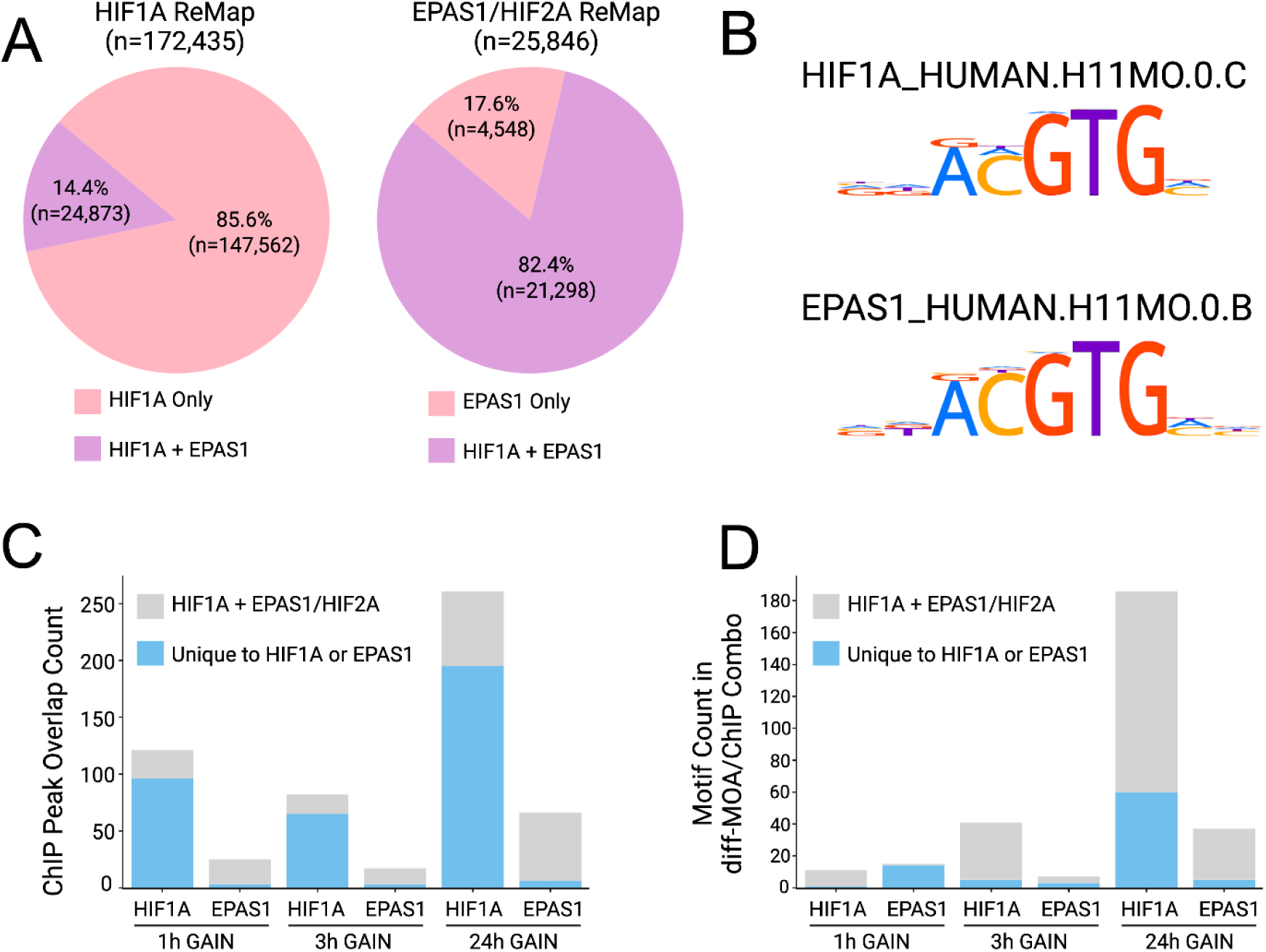
Comparison of EPAS1/HIF2A and HIF1A ChIP-seq Overlap and Motif Intersections. **(A)** Pie charts showing the overlap between the HIF1A ReMap ChIP-seq data set and the EPAS1/HIF2A ChIP-seq data set. The pink sections represent peaks unique to each ChIP-seq dataset, while the lavender sections represent overlapping peaks. For HIF1A, 85.6% of its peaks are unique, while 14.4% are shared. For EPAS1/HIF2A, only 17.6% of its peaks are unique, while 82.4% are shared. **(B)** Motif logos from HOCOMOCOv11 for both HIF1A and EPAS1/HIF2A. **(C)** ChIP-seq peak intersections: The intersection of each set of gain diff-MOA peaks by hour with the HIF1A and EPAS1/HIF2A ChIP-seq datasets. Bars are color-coded to indicate the proportion that is unique to the ChIP-seq dataset (blue) and the proportion that is shared or overlapping (grey). **(D)** Motif intersections: The intersection of each set of gain diff-MOA peaks by hour with the HIF1A and EPAS1/HIF2A motifs found via the MEME suite. Bars are color-coded to indicate the proportion of regions exclusive to the motif (blue) and the proportion of regions with overlapping motifs (grey).

## References

Allan, K. C., Hu, L. R., Scavuzzo, M. A., Morton, A. R., Gevorgyan, A. S., Cohn, E. F., Clayton, B. L. L., Bederman, I. R., Hung, S., Bartels, C. F., Madhavan, M., & Tesar, P. J. (2021). Non-canonical targets of HIF1a impair oligodendrocyte progenitor cell function. Cell Stem Cell, 28(2), 257–272.e11.

Amemiya, H. M., Kundaje, A., & Boyle, A. P. (2019). The ENCODE Blacklist: Identification of Problematic Regions of the Genome. Scientific Reports, 9(1), 9354.

An integrated encyclopedia of DNA elements in the human genome. (2012). Nature, 489(7414), 57–74.

Awad, H., Nolette, N., Hinton, M., & Dakshinamurti, S. (2014). AMPK and FoxO1 regulate catalase expression in hypoxic pulmonary arterial smooth muscle. Pediatric Pulmonology, 49(9), 885–897.

Baek, S., & Sung, M.-H. (2016). Genome-scale analysis of cell-specific regulatory codes using nuclear enzymes. *Methods in Molecular Biology (Clifton*, N.J.), 1418, 225–240.

Bae, T., Hallis, S. P., & Kwak, M.-K. (2024). Hypoxia, oxidative stress, and the interplay of HIFs and NRF2 signaling in cancer. Experimental & Molecular Medicine, 56(3), 501–514.

Bailey, T. L., Johnson, J., Grant, C. E., & Noble, W. S. (2015). The MEME suite. Nucleic Acids Research, 43(W1), W39–W49.

Bartoszewski, R., Moszyńska, A., Serocki, M., Cabaj, A., Polten, A., Ochocka, R., Dell’Italia, L., Bartoszewska, S., Króliczewski, J., Dąbrowski, M., & Collawn, J. F. (2019). Primary endothelial cell-specific regulation of hypoxia-inducible factor (HIF)-1 and HIF-2 and their target gene expression profiles during hypoxia. FASEB Journal: Official Publication of the Federation of American Societies for Experimental Biology, 33(7), 7929–7941.

Batie, M., Frost, J., Frost, M., Wilson, J. W., Schofield, P., & Rocha, S. (2019). Hypoxia induces rapid changes to histone methylation and reprograms chromatin. *Science (New York*, N.Y.), 363(6432), 1222–1226.

Batie, M., & Rocha, S. (2020). Gene transcription and chromatin regulation in hypoxia. Biochemical Society Transactions, 48(3), 1121–1128.

Benoit, J., Sheikhbahaei, M. K., & Dennis, J. (2021). Chromatin dynamics: Nucleosome occupancy and sensitivity as determinants of gene expression and cell fate. Journal of Cancer Biology, 2(2), 51–55.

Brahma, S., & Henikoff, S. (2019). RSC-associated subnucleosomes define MNase-sensitive promoters in yeast. Molecular Cell, 73(2), 238–249.e3.

Carone, B. R. (2025). MNase-seq to identify genome-wide DNA-protein interactions. *Methods in Molecular Biology (Clifton*, N.J.), 2866, 99–109.

Casey, B. H., Kollipara, R. K., Pozo, K., & Johnson, J. E. (2018). Intrinsic DNA binding properties demonstrated for lineage-specifying basic helix-loop-helix transcription factors. Genome Research, 28(4), 484–496.

Chen, E. Y., Tan, C. M., Kou, Y., Duan, Q., Wang, Z., Meirelles, G. V., Clark, N. R., & Ma’ayan, A. (2013). Enrichr: interactive and collaborative HTML5 gene list enrichment analysis tool. BMC Bioinformatics, 14, 128.

Chi, J.-T., Wang, Z., Nuyten, D. S. A., Rodriguez, E. H., Schaner, M. E., Salim, A., Wang, Y., Kristensen, G. B., Helland, A., Børresen-Dale, A.-L., Giaccia, A., Longaker, M. T., Hastie, T., Yang, G. P., van de Vijver, M. J., & Brown, P. O. (2006). Gene expression programs in response to hypoxia: cell type specificity and prognostic significance in human cancers. PLoS Medicine, 3(3), e47.

Chomczynski, P. (1993). A reagent for the single-step simultaneous isolation of RNA, DNA and proteins from cell and tissue samples. BioTechniques, 15(3), 532–534, 536–537.

Cirillo, F., Resmini, G., Ghiroldi, A., Piccoli, M., Bergante, S., Tettamanti, G., & Anastasia, L. (2017). Activation of the hypoxia-inducible factor 1α promotes myogenesis through the noncanonical Wnt pathway, leading to hypertrophic myotubes. FASEB Journal: Official Publication of the Federation of American Societies for Experimental Biology, 31(5), 2146–2156.

Crusoe, M. R., Alameldin, H. F., Awad, S., Boucher, E., Caldwell, A., Cartwright, R., Charbonneau, A., Constantinides, B., Edvenson, G., Fay, S., Fenton, J., Fenzl, T., Fish, J., Garcia-Gutierrez, L., Garland, P., Gluck, J., González, I., Guermond, S., Guo, J., … Brown, C. T. (2015). The khmer software package: enabling efficient nucleotide sequence analysis. F1000Research, 4, 900.

ENCODE Project Consortium, Moore, J. E., Purcaro, M. J., Pratt, H. E., Epstein, C. B., Shoresh, N., Adrian, J., Kawli, T., Davis, C. A., Dobin, A., Kaul, R., Halow, J., Van Nostrand, E. L., Freese, P., Gorkin, D. U., Shen, Y., He, Y., Mackiewicz, M., Pauli-Behn, F., … Weng, Z. (2020). Expanded encyclopaedias of DNA elements in the human and mouse genomes. Nature, 583(7818), 699–710.

Engelhorn, J., Snodgrass, S. J., Kok, A., Seetharam, A. S., Schneider, M., Kiwit, T., Singh, A., Banf, M., Doan, D. T. H., Khaipho-Burch, M., Runcie, D. E., Sánchez-Camargo, V. A., Bader, R., Vladimir Torres-Rodriguez, J., Sun, G., Stam, M., Fiorani, F., Beier, S., Schnable, J. C., … Hartwig, T. (2025). Genetic variation at transcription factor binding sites largely explains phenotypic heritability in maize. Nature Genetics, 57(9), 2313–2322.

Esnault, C., Magat, T., García-Oliver, E., & Andrau, J.-C. (2021). Analyses of promoter, enhancer, and nucleosome organization in mammalian cells by MNase-Seq. *Methods in Molecular Biology (Clifton*, N.J.), 2351, 93–104.

Frankish, A., Carbonell-Sala, S., Diekhans, M., Jungreis, I., Loveland, J. E., Mudge, J. M., Sisu, C., Wright, J. C., Arnan, C., Barnes, I., Banerjee, A., Bennett, R., Berry, A., Bignell, A., Boix, C., Calvet, F., Cerdán-Vélez, D., Cunningham, F., Davidson, C., … Flicek, P. (2023). GENCODE: reference annotation for the human and mouse genomes in 2023. Nucleic Acids Research, 51(D1), D942–D949.

Hammal, F., de Langen, P., Bergon, A., Lopez, F., & Ballester, B. (2022). ReMap 2022: a database of Human, Mouse, Drosophila and Arabidopsis regulatory regions from an integrative analysis of DNA-binding sequencing experiments. Nucleic Acids Research, 50(D1), D316–D325.

Hernansanz-Agustín, P., Choya-Foces, C., Enríquez, J. A., & Martínez-Ruiz, A. (2021). Na+ controls hypoxic signalling by the mitochondrial respiratory chain. Free Radical Biology & Medicine, 165, 18.

Hu, G., Grover, C. E., Vera, D. L., Lung, P.-Y., Girimurugan, S. B., Miller, E. R., Conover, J. L., Ou, S., Xiong, X., Zhu, D., Li, D., Gallagher, J. P., Udall, J. A., Sui, X., Zhang, J., Bass, H. W., & Wendel, J. F. (2024). Evolutionary dynamics of chromatin structure and duplicate gene expression in diploid and allopolyploid cotton. Molecular Biology and Evolution, 41(5). 10.1093/molbev/msae095

Kakani, P., Dhamdhere, S. G., Pant, D., Joshi, R., Mishra, S., Pandey, A., Notani, D., & Shukla, S. (2025). Hypoxia-induced CTCF mediates alternative splicing via coupling chromatin looping and RNA Pol II pause to promote EMT in breast cancer. Cell Reports, 44(2), 115267.

Kaluz, S., Kaluzová, M., & Stanbridge, E. J. (2008). Regulation of gene expression by hypoxia: integration of the HIF-transduced hypoxic signal at the hypoxia-responsive element. Clinica Chimica Acta; International Journal of Clinical Chemistry, 395(1-2), 6–13.

Kent, W. J., Zweig, A. S., Barber, G., Hinrichs, A. S., & Karolchik, D. (2010). BigWig and BigBed: enabling browsing of large distributed datasets. *Bioinformatics (Oxford*, England), 26(17), 2204–2207.

Kindrick, J. D., & Mole, D. R. (2020). Hypoxic regulation of gene transcription and chromatin: Cause and effect. International Journal of Molecular Sciences, 21(21), 8320.

Koltsova, S. V., Shilov, B., Birulina, J. G., Akimova, O. A., Haloui, M., Kapilevich, L. V., Gusakova, S. V., Tremblay, J., Hamet, P., & Orlov, S. N. (2014). Transcriptomic changes triggered by hypoxia: evidence for HIF-1α-independent, [Na+]i/[K+]i-mediated, excitation-transcription coupling. PloS One, 9(11), e110597.

Kulakovskiy, I. V., Vorontsov, I. E., Yevshin, I. S., Sharipov, R. N., Fedorova, A. D., Rumynskiy, E. I., Medvedeva, Y. A., Magana-Mora, A., Bajic, V. B., Papatsenko, D. A., Kolpakov, F. A., & Makeev, V. J. (2018). HOCOMOCO: towards a complete collection of transcription factor binding models for human and mouse via large-scale ChIP-Seq analysis. Nucleic Acids Research, 46(D1), D252–D259.

Kuleshov, M. V., Jones, M. R., Rouillard, A. D., Fernandez, N. F., Duan, Q., Wang, Z., Koplev, S., Jenkins, S. L., Jagodnik, K. M., Lachmann, A., McDermott, M. G., Monteiro, C. D., Gundersen, G. W., & Ma’ayan, A. (2016). Enrichr: a comprehensive gene set enrichment analysis web server 2016 update. Nucleic Acids Research, 44(W1), W90–W97.

Landt, S. G., Marinov, G. K., Kundaje, A., Kheradpour, P., Pauli, F., Batzoglou, S., Bernstein, B. E., Bickel, P., Brown, J. B., Cayting, P., Chen, Y., DeSalvo, G., Epstein, C., Fisher-Aylor, K. I., Euskirchen, G., Gerstein, M., Gertz, J., Hartemink, A. J., Hoffman, M. M., … Snyder, M. (2012). ChIP-seq guidelines and practices of the ENCODE and modENCODE consortia. Genome Research, 22(9), 1813–1831.

Lee, P., Chandel, N. S., & Simon, M. C. (2020). Cellular adaptation to hypoxia through hypoxia inducible factors and beyond. Nature Reviews. Molecular Cell Biology, 21(5), 268–283.

Liang, G.-P., Su, Y.-Y., Chen, J., Yang, Z.-C., Liu, Y.-S., & Luo, X.-D. (2009). Analysis of the early adaptive response of endothelial cells to hypoxia via a long serial analysis of gene expression. Biochemical and Biophysical Research Communications, 384(4), 415–419.

Li, H., Handsaker, B., Wysoker, A., Fennell, T., Ruan, J., Homer, N., Marth, G., Abecasis, G., Durbin, R., & 1000 Genome Project Data Processing Subgroup. (2009). The Sequence Alignment/Map format and SAMtools. Bioinformatics, 25(16), 2078–2079.

Li, Q., Brown, J. B., Huang, H., & Bickel, P. J. (2011). Measuring reproducibility of high-throughput experiments. The Annals of Applied Statistics, 5(3), 1752–1779.

Liu, O. H.-F., Kiema, M., Beter, M., Ylä-Herttuala, S., Laakkonen, J. P., & Kaikkonen, M. U. (2020). Hypoxia-mediated regulation of histone demethylases affects angiogenesis-associated functions in endothelial cells. *Arteriosclerosis*, Thrombosis, and Vascular Biology, 40(11), 2665–2677.

Love, M. I., Huber, W., & Anders, S. (2014). Moderated estimation of fold change and dispersion for RNA-seq data with DESeq2. Genome Biology, 15(12), 550.

Luo, K., Zhong, J., Safi, A., Hong, L. K., Tewari, A. K., Song, L., Reddy, T. E., Ma, L., Crawford, G. E., & Hartemink, A. J. (2022). Profiling the quantitative occupancy of myriad transcription factors across conditions by modeling chromatin accessibility data. Genome Research, 32(6), 1183–1198.

Magoč, T., & Salzberg, S. L. (2011). FLASH: fast length adjustment of short reads to improve genome assemblies. Bioinformatics, 27(21), 2957–2963.

Martin, M. (2011). Cutadapt removes adapter sequences from high-throughput sequencing reads. EMBnet.journal, 17(1), 10–12.

Mole, D. R., Blancher, C., Copley, R. R., Pollard, P. J., Gleadle, J. M., Ragoussis, J., & Ratcliffe, P. J. (2009). Genome-wide association of hypoxia-inducible factor (HIF)-1alpha and HIF-2alpha DNA binding with expression profiling of hypoxia-inducible transcripts. The Journal of Biological Chemistry, 284(25), 16767–16775.

Mootha, V. K., Lindgren, C. M., Eriksson, K.-F., Subramanian, A., Sihag, S., Lehar, J., Puigserver, P., Carlsson, E., Ridderstråle, M., Laurila, E., Houstis, N., Daly, M. J., Patterson, N., Mesirov, J. P., Golub, T. R., Tamayo, P., Spiegelman, B., Lander, E. S., Hirschhorn, J. N., … Groop, L. C. (2003). PGC-1α-responsive genes involved in oxidative phosphorylation are coordinately downregulated in human diabetes. Nature Genetics, 34(3), 267–273.

Niskanen, H., Tuszynska, I., Zaborowski, R., Heinäniemi, M., Ylä-Herttuala, S., Wilczynski, B., & Kaikkonen, M. U. (2018). Endothelial cell differentiation is encompassed by changes in long range interactions between inactive chromatin regions. Nucleic Acids Research, 46(4), 1724–1740.

Ohnishi, S., Maehara, O., Nakagawa, K., Kameya, A., Otaki, K., Fujita, H., Higashi, R., Takagi, K., Asaka, M., Sakamoto, N., Kobayashi, M., & Takeda, H. (2013). hypoxia-inducible factors activate CD133 promoter through ETS family transcription factors. PloS One, 8(6), e66255.

Ohno, M., Ando, T., Priest, D. G., Kumar, V., Yoshida, Y., & Taniguchi, Y. (2019). Sub-nucleosomal genome structure reveals distinct nucleosome folding motifs. Cell, 176(3), 520–534.e25.

Parvathaneni, R. K., Bertolini, E., Shamimuzzaman, M., Vera, D. L., Lung, P.-Y., Rice, B. R., Zhang, J., Brown, P. J., Lipka, A. E., Bass, H. W., & Eveland, A. L. (2020). The regulatory landscape of early maize inflorescence development. Genome Biology, 21(1), 165.

Payne, S., Neal, A., & De Val, S. (2024). Transcription factors regulating vasculogenesis and angiogenesis. Developmental Dynamics: An Official Publication of the American Association of Anatomists, 253(1), 28–58.

Pullamsetti, S. S., Mamazhakypov, A., Weissmann, N., Seeger, W., & Savai, R. (2020). Hypoxia-inducible factor signaling in pulmonary hypertension. The Journal of Clinical Investigation, 130(11), 5638–5651.

Quinlan, A. R., & Hall, I. M. (2010). BEDTools: a flexible suite of utilities for comparing genomic features. Bioinformatics, 26(6), 841–842.

Ramírez, F., Dündar, F., Diehl, S., Grüning, B. A., & Manke, T. (2014). deepTools: a flexible platform for exploring deep-sequencing data. Nucleic Acids Research, 42(Web Server issue), W187–W191.

Ren, G., Jin, W., Cui, K., Rodrigez, J., Hu, G., Zhang, Z., Larson, D. R., & Zhao, K. (2017). CTCF-mediated enhancer-promoter interaction is a critical regulator of cell-to-cell variation of gene expression. Molecular Cell, 67(6), 1049–1058.e6.

Rodgers-Melnick, E., Vera, D. L., Bass, H. W., & Buckler, E. S. (2016). Open chromatin reveals the functional maize genome. Proceedings of the National Academy of Sciences of the United States of America, 113(22), E3177–E3184.

Sabo, P. J., Hawrylycz, M., Wallace, J. C., Humbert, R., Yu, M., Shafer, A., Kawamoto, J., Hall, R., Mack, J., Dorschner, M. O., McArthur, M., & Stamatoyannopoulos, J. A. (2004). Discovery of functional noncoding elements by digital analysis of chromatin structure. Proceedings of the National Academy of Sciences of the United States of America, 101(48), 16837–16842.

Sabo, P. J., Kuehn, M. S., Thurman, R., Johnson, B. E., Johnson, E. M., Cao, H., Yu, M., Rosenzweig, E., Goldy, J., Haydock, A., Weaver, M., Shafer, A., Lee, K., Neri, F., Humbert, R., Singer, M. A., Richmond, T. A., Dorschner, M. O., McArthur, M., … Stamatoyannopoulos, J. A. (2006). Genome-scale mapping of DNase I sensitivity in vivo using tiling DNA microarrays. Nature Methods, 3(7), 511–518.

Savadel, S. D., Hartwig, T., Turpin, Z. M., Vera, D. L., Lung, P.-Y., Sui, X., Blank, M., Frommer, W. B., Dennis, J. H., Zhang, J., & Bass, H. W. (2021). The native cistrome and sequence motif families of the maize ear. PLoS Genetics, 17(8), e1009689.

Scheurer, S. B., Raybak, J. N., Rösli, C., Neri, D., & Elia, G. (2004). Modulation of gene expression by hypoxia in human umbilical cord vein endothelial cells: A transcriptomic and proteomic study (vol. 4, Issue 6, pp. 1737–1760). *Proteomics*, *4*(9), 2822–2822.

Schmitges, F. W., Radovani, E., Najafabadi, H. S., Barazandeh, M., Campitelli, L. F., Yin, Y., Jolma, A., Zhong, G., Guo, H., Kanagalingam, T., Dai, W. F., Taipale, J., Emili, A., Greenblatt, J. F., & Hughes, T. R. (2016). Multiparameter functional diversity of human C2H2 zinc finger proteins. Genome Research, 26(12), 1742–1752.

Schödel, J., Oikonomopoulos, S., Ragoussis, J., Pugh, C. W., Ratcliffe, P. J., & Mole, D. R. (2011). High-resolution genome-wide mapping of HIF-binding sites by ChIP-seq. Blood, 117(23), e207–e217.

Smythies, J. A., Sun, M., Masson, N., Salama, R., Simpson, P. D., Murray, E., Neumann, V., Cockman, M. E., Choudhry, H., Ratcliffe, P. J., & Mole, D. R. (2019). Inherent DNA-binding specificities of the HIF-1α and HIF-2α transcription factors in chromatin. EMBO Reports, 20(1), e46401.

Subramanian, A., Tamayo, P., Mootha, V. K., Mukherjee, S., Ebert, B. L., Gillette, M. A., Paulovich, A., Pomeroy, S. L., Golub, T. R., Lander, E. S., & Mesirov, J. P. (2005). Gene set enrichment analysis: a knowledge-based approach for interpreting genome-wide expression profiles. Proceedings of the National Academy of Sciences of the United States of America, 102(43), 15545–15550.

Suzuki, N., Vojnovic, N., Lee, K.-L., Yang, H., Gradin, K., & Poellinger, L. (2018). HIF-dependent and reversible nucleosome disassembly in hypoxia-inducible gene promoters. Experimental Cell Research, 366(2), 181–191.

Vasimuddin, M., Misra, S., Li, H., & Aluru, S. (2019). Efficient Architecture-Aware Acceleration of BWA-MEM for Multicore Systems. 2019 IEEE International Parallel and Distributed Processing Symposium (IPDPS), 314–324.

Vierstra, J., Lazar, J., Sandstrom, R., Halow, J., Lee, K., Bates, D., Diegel, M., Dunn, D., Neri, F., Haugen, E., Rynes, E., Reynolds, A., Nelson, J., Johnson, A., Frerker, M., Buckley, M., Kaul, R., Meuleman, W., & Stamatoyannopoulos, J. A. (2020). Global reference mapping of human transcription factor footprints. Nature, 583(7818), 729–736.

Villar, D., Ortiz-Barahona, A., Gómez-Maldonado, L., Pescador, N., Sánchez-Cabo, F., Hackl, H., Rodriguez, B. A. T., Trajanoski, Z., Dopazo, A., Huang, T. H. M., Yan, P. S., & Del Peso, L. (2012). Cooperativity of stress-responsive transcription factors in core hypoxia-inducible factor binding regions. PloS One, 7(9), e45708.

Wang, Q., Li, M., Wu, T., Zhan, L., Li, L., Chen, M., Xie, W., Xie, Z., Hu, E., Xu, S., & Yu, G. (2022). Exploring epigenomic datasets by ChIPseeker. Current Protocols, 2(10), e585.

Wang, S., Chen, J., Garcia, S. P., Liang, X., Zhang, F., Yan, P., Yu, H., Wei, W., Li, Z., Wang, J., Le, H., Han, Z., Luo, X., Day, D. S., Stevens, S. M., Zhang, Y., Park, P. J., Liu, Z.-J., Sun, K., … Zhang, B. (2019). A dynamic and integrated epigenetic program at distal regions orchestrates transcriptional responses to VEGFA. Genome Research, 29(2), 193–207.

Weckmann, K., Diefenthäler, P., Baeken, M. W., Yusifli, K., Turck, C. W., Asara, J. M., Behl, C., & Hajieva, P. (2018). Metabolomics profiling reveals differential adaptation of major energy metabolism pathways associated with autophagy upon oxygen and glucose reduction. Scientific Reports, 8(1), 2337.

West, J. B. (2017). Physiological effects of chronic hypoxia. The New England Journal of Medicine, 376(20), 1965–1971.

Wickham, H. (n.d.). *ggplot2*. Springer New York.

Wollenick, K., Hu, J., Kristiansen, G., Schraml, P., Rehrauer, H., Berchner-Pfannschmidt, U., Fandrey, J., Wenger, R. H., & Stiehl, D. P. (2012). Synthetic transactivation screening reveals ETV4 as broad coactivator of hypoxia-inducible factor signaling. Nucleic Acids Research, 40(5), 1928–1943.

Wu, D., Dasgupta, A., Read, A. D., Bentley, R. E. T., Motamed, M., Chen, K.-H., Al-Qazazi, R., Mewburn, J. D., Dunham-Snary, K. J., Alizadeh, E., Tian, L., & Archer, S. L. (2021). Oxygen sensing, mitochondrial biology and experimental therapeutics for pulmonary hypertension and cancer. Free Radical Biology & Medicine, 170, 150–178.

Xie, Z., Bailey, A., Kuleshov, M. V., Clarke, D. J. B., Evangelista, J. E., Jenkins, S. L., Lachmann, A., Wojciechowicz, M. L., Kropiwnicki, E., Jagodnik, K. M., Jeon, M., & Ma’ayan, A. (2021). Gene Set Knowledge Discovery with Enrichr. Current Protocols, 1(3), e90.

Yadav, S., Kalra, N., Ganju, L., & Singh, M. (2017). Activator protein-1 (AP-1): a bridge between life and death in lung epithelial (A549) cells under hypoxia. Molecular and Cellular Biochemistry, 436(1-2), 99–110.

Yan, F., Powell, D. R., Curtis, D. J., & Wong, N. C. (2020). From reads to insight: a hitchhiker’s guide to ATAC-seq data analysis. Genome Biology, 21(1), 22.

Yfantis, A., Mylonis, I., Chachami, G., Nikolaidis, M., Amoutzias, G. D., Paraskeva, E., & Simos, G. (2023). Transcriptional response to hypoxia: The role of HIF-1-associated co-regulators. Cells (Basel, Switzerland), 12(5). 10.3390/cells12050798

Yoshimoto, S., Tanaka, F., Morita, H., Hiraki, A., & Hashimoto, S. (2019). Hypoxia-induced HIF-1α and ZEB1 are critical for the malignant transformation of ameloblastoma via TGF-β-dependent EMT. Cancer Medicine, 8(18), 7822–7832.

Yu, G., Wang, L.-G., & He, Q.-Y. (2015). ChIPseeker: an R/Bioconductor package for ChIP peak annotation, comparison and visualization. *Bioinformatics (Oxford*, England), 31(14), 2382–2383.

Zhang, B., Day, D. S., Ho, J. W., Song, L., Cao, J., Christodoulou, D., Seidman, J. G., Crawford, G. E., Park, P. J., & Pu, W. T. (2013). A dynamic H3K27ac signature identifies VEGFA-stimulated endothelial enhancers and requires EP300 activity. Genome Research, 23(6), 917–927.

Zhang, B., Wu, J., Cai, Y., Luo, M., Wang, B., & Gu, Y. (2019). TCF7L1 indicates prognosis and promotes proliferation through activation of Keap1/NRF2 in gastric cancer. Acta Biochimica et Biophysica Sinica, 51(4), 375–385.

Zhang, Y., Liu, T., Meyer, C. A., Eeckhoute, J., Johnson, D. S., Bernstein, B. E., Nusbaum, C., Myers, R. M., Brown, M., Li, W., & Liu, X. S. (2008). Model-based analysis of ChIP-Seq (MACS). Genome Biology, 9(9), R137.

Zheng, J., Kim, S.-J., Saeidi, S., Kim, S. H., Fang, X., Lee, Y.-H., Guillen-Quispe, Y. N., Ngo, H. K. C., Kim, D.-H., Kim, D., & Surh, Y.-J. (2023). Overactivated NRF2 induces pseudohypoxia in hepatocellular carcinoma by stabilizing HIF-1α. Free Radical Biology & Medicine, 194, 347–356.

