## Supplementary figures and images for "The cistrome response to hypoxia in human umbilical vein endothelial cells"

### logo1.png

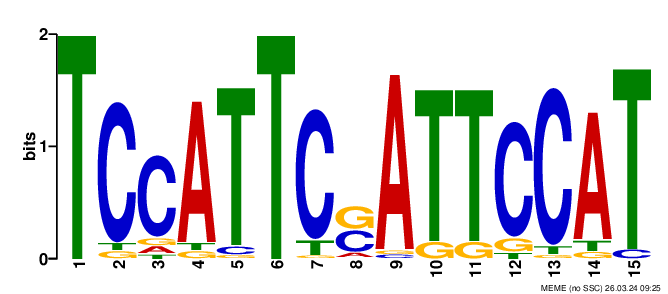

### logo1.png

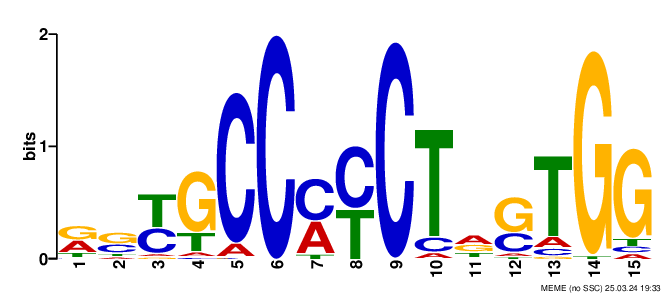

### logo2.png

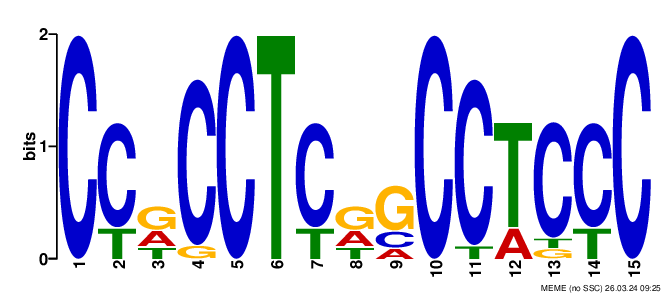

### logo2.png

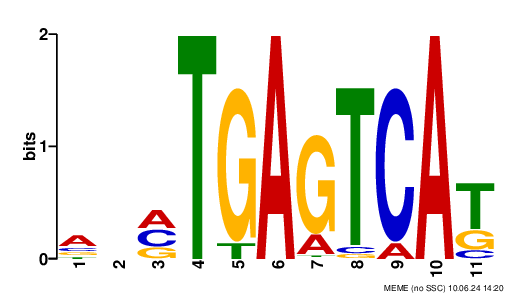

### logo4.png

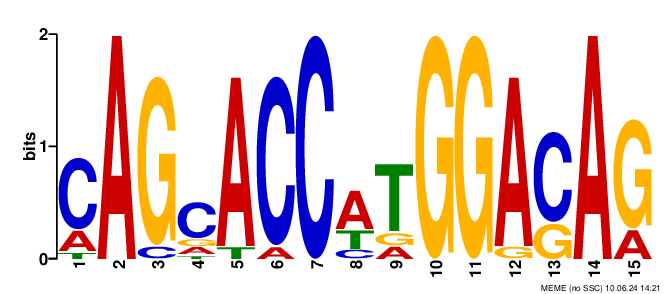

### logo_rc1.png

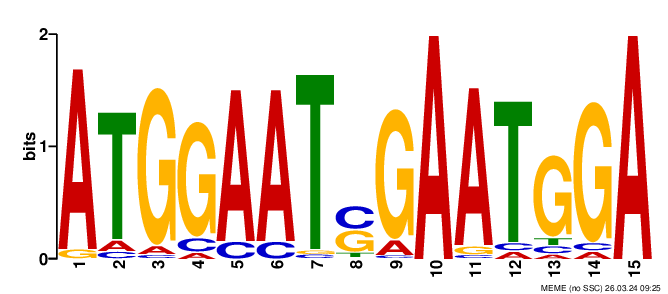

### logo_rc1.png

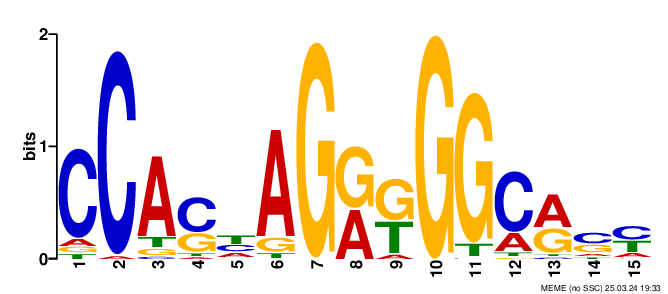

### logo_rc1.png

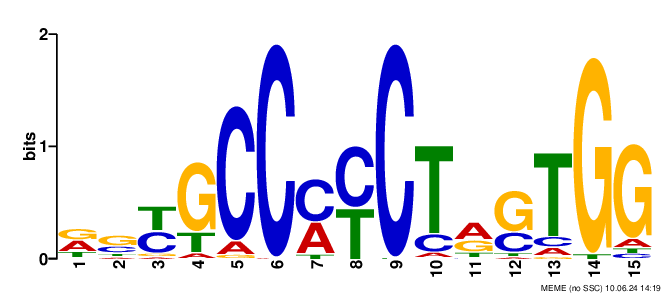

### logo_rc2.png

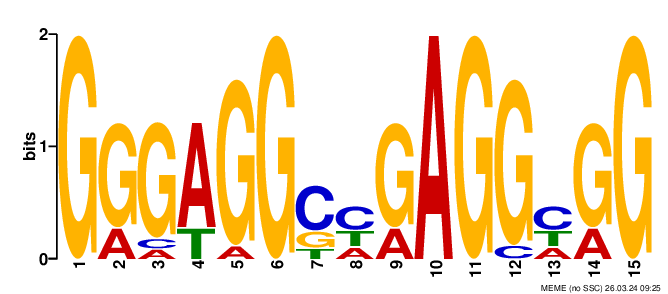

### logo_rc2.png

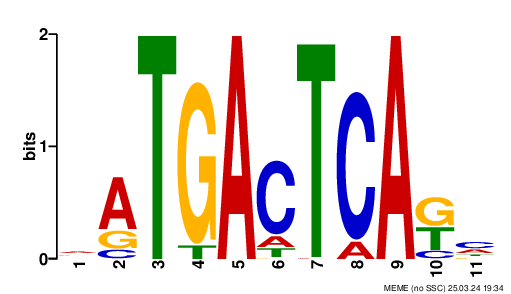

### logo_rc2.png

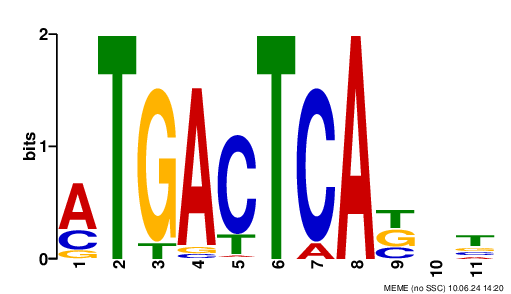

### logo_rc3.png

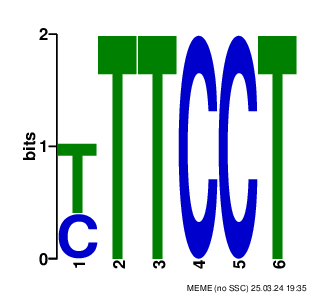

### logo_rc3.png

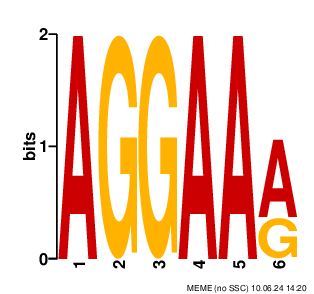

### logo_rc4.png

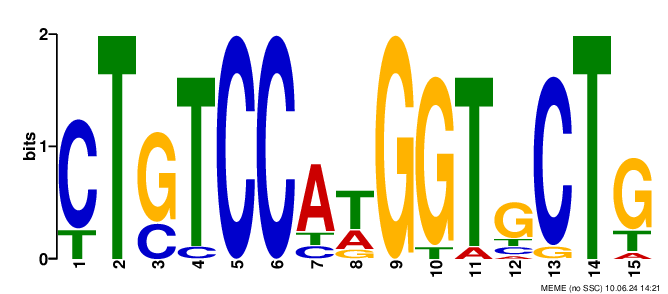
